# Sleep spindles stabilize human thalamocortical networks, inhibiting pathological disruptions

**DOI:** 10.64898/2026.09.13.751011

**Authors:** Kassem Jaber, Alyssa Ho, Hongyi Ye, Pilar Bosque-Varela, Giovanna Aiello, Tamir Avigdor, Krishma Shenoy, Praveen Ramani, Lara Wadi, John Thomas, Petr Klimeš, R. Mark Richardson, Derek Southwell, Peter N. Hadar, Sydney S. Cash, Pariya Salami, David Carlson, Rina Zelmann, Birgit Frauscher

**Affiliations:** Analytical Neurophysiology Lab, Department of Neurology, Duke University Medical Center, North Carolina, U.S.A; Department of Biomedical Engineering, Duke Pratt School of Engineering, Durham, North Carolina U.S.A; Department of Neurology and Epilepsy Center, The Second Affiliated Hospital, Zhejiang University School of Medicine, Hangzhou, China; Department of Neurology, Neurocritical Care and Neurorehabilitation, Christian Doppler University Hospital, Centre for Cognitive Neuroscience, Member of the European Reference Network EpiCARE, Paracelsus Medical University of Salzburg, 5020 Salzburg, Austria; ETH Zurich, Department of Health Sciences and Technology, Institute for Neuroscience, ETH Zürich, 8092 Zürich, Switzerland; Montreal Neurological Institute and Hospital, McGill University, Montréal, Québec H3A 2B4, Canada; Department of Neurology, Mass General Brigham and Harvard Medical School, Boston, MA, USA; Center for Neurotechnology and Neurorecovery, Massachusetts General Hospital, Boston, MA, USA; Department of Pediatrics, University of Arkansas for Medical Sciences, Arkansas, USA; Department of Neurology, Duke University Medical Center, North Carolina, USA; Department of Electrical and Computer Engineering Technology, Rochester Institute of Technology, NY, USA; Department of Neurosurgery, Mass General Brigham and Harvard Medical School, Boston, MA, USA; Department of Neurosurgery, Duke University Medical Center, Durham, NC, USA; Duke University Institute of Statistics and Decision Sciences, Durham, North Carolina, USA; Department of Computer Science, Department of Civil and Environmental Engineering, Duke University, Durham, North Carolina, USA

## Abstract

The thalamus coordinates sleep-dependent brain function including memory consolidation, sensory gating, and network stability through spindle oscillations. Thalamocortical circuits are also affected in many neurological and neuropsychiatric disorders. Yet how human thalamocortical circuits respond to pathological disruptions remain poorly understood. Here, we leveraged interictal epileptiform discharges (IEDs) as spontaneous perturbations to probe spindle-generating circuits and thalamocortical dynamics in vivo. Using multi-night intracranial recordings spanning multiple thalamic nuclei and cortical regions in 55 individuals with epilepsy, we investigated interactions between sleep spindles and epileptic activity across timescales from milliseconds to days. Sleep spindles were associated with IED suppression, whereas IEDs increased subsequent spindle probability, revealing bidirectional interactions between physiological and pathological activity. Nights with lower IEDs corresponded to increased spindle occurrence, longer duration, and faster frequency, reflecting a more stable thalamocortical state. These findings propose sleep spindles as potential regulators of thalamocortical networks that actively respond to and regulate pathological activity.

## INTRODUCTION

The thalamus serves as a central hub in the brain, integrating and relaying information across cortical regions. It plays a dynamic role in coordinating cognitive function^1^, sensory processing^2^, sleep regulation^3,4^ and memory consolidation^5,6^ through thalamocortical interactions^7^. These interactions are central to the generation of sleep spindles and the regulation of cortical excitability during non-rapid-eye movement (NREM) sleep^8–12^. Sleep spindles are physiological NREM oscillations (10-16Hz) essential for cognitive function, memory consolidation, and sleep physiology^13^. They are thought to be generated by bidirectional interactions between the thalamic reticular nucleus and thalamocortical relay neurons^14^. Important for brain health, sleep spindles are a recognized electrographic marker of stable sleep^15^.

In epilepsy, the thalamus is increasingly recognized as a key player in both seizure propagation and maintenance^16–18^. However, most of our understanding of how the thalamus functions is derived from animal studies^14,19–22^. Direct evidence in humans remains scarce, as the thalamus is a deep structure which activity cannot be detected using standard scalp electroencephalogram (EEG). Recently, however, the thalamus has emerged as a key-target for thalamic neuromodulation strategies such as deep-brain stimulation (DBS) and responsive neurostimulation (RNS), and consequently invasive intracranial stereo-EEG (sEEG) investigations for drug-resistant epilepsy^23,24^. During presurgical investigation, sEEG recordings are used to interrogate the human brain, recently including thalamic nuclei as nodes in patient-specific seizure networks to identify targets for neuromodulation. This sEEG data provides a unique opportunity to investigate *in-vivo* human thalamocortical interactions.

Sleep and epilepsy have complex bidirectional interactions: sleep instability significantly affects people with epilepsy^25–27^, regardless of whether their seizures are sleep related^28^. Inducing spindles during sleep were shown to increase NREM durations in mice with more NREM-REM transitions^29^ while greater spindle activity in humans were associated with increased resistance to sleep disruption^15,30^. Therefore, sleep spindles have been proposed to stabilize sleep^10,15^ by filtering sensory inputs through thalamocortical mechanisms involving inhibitory reticular activity and rebound firing in thalamocortical relay cells^15,29,31^.

While spindles have been positively associated with cognition^32^, interictal epileptiform discharges (IEDs), defined as brief (<200ms) EEG transients generated by epileptogenic tissue^33^, were shown to impair cognition^34–36^.Therefore, IEDs likely have a complex pathophysiological interaction with spindle-generating circuits^37–41^. For example, hippocampal IEDs were shown to induce spindles in the prefrontal cortex^37,42^ and IED-spindle coupling was associated with poor memory performance^42^, likely suggesting that IEDs ‘hijack’ the reticular-thalamic circuit, reflected by the time-locked facilitation of spindles^37^. These associations of spindles with physiological memory processing, sleep stability, and pathological epileptic activity introduce a paradox that remains to be fully understood.

*We hypothesize that the inverse relation between sleep stability and pathological brain excitability is mediated by thalamocortical dynamics, and that thalamic sleep spindles reflect a shift toward a physiological steady-state, making the thalamocortical network less susceptible to pathological engagement.* We tested this hypothesis by leveraging IEDs as spontaneous perturbations to investigate how spindle-generating circuits interact with epileptic networks and to understand the role of these circuits in regulating network stability. A clearer mechanistic understanding of thalamocortical circuit pathophysiology is needed to improve pharmacological management of epilepsy and to optimize hypothesis-driven neuromodulation strategies for drug-resistant epilepsy. This would provide the first evidence of a control system linking sleep stability and brain excitability, with implications beyond epilepsy and across neurological and neuropsychiatric domains.

Here, we investigated how sleep stability reshapes epileptic activity. Specifically, we tested the hypothesis that spindle activity protects against pathological disturbances. We did this by characterizing interactions between sleep spindles and IEDs across long (i.e., multi-night) and short (i.e., hundred-millisecond) timescales in drug-resistant focal epilepsy. The seizure-onset zone (SOZ) – a marker of epileptogenic tissue and defined as the subset of sEEG channels exhibiting the earliest changes when seizures start^43^ – was used to delineate the diseased tissue to investigate IED activity and its interaction with thalamic sleep spindles. IEDs have been shown to fluctuate at patient-specific circadian rhythms^44,45^. Therefore, for long timescale analysis, we selected nights with minimum and maximum IED rates for each patient to capture the largest available contrast. More specifically, we quantified thalamo-epileptic interactions by (i) contrasting thalamic spindle structure across nights of minimum and maximum IED rates in the SOZ; (ii) time-locking thalamic spindles to measure their influence on IEDs in the SOZ; and (iii) exploring thalamic nuclei involvement in these dynamics within this cohort. If stable sleep is protective against epileptic activity, these results may identify sleep stability as a clinically relevant brain state for state-aware neuromodulation to treat people with drug-resistant epilepsy.

## RESULTS

### Dataset Demographics

We screened 58 consecutive patients from Duke University Hospital (DUH; 2023-2024), and Massachusetts General Hospital (MGH; 2020-2024), who underwent phase II presurgical investigation with at least one SEEG electrode sampling the thalamus. Fifty-five patients met the selection criteria described in **Methods** and were included in the analysis (**Figure S1**). Of the 55 included patients, 27 (49%) were female, mean age 30.2 ± 15.2 years. Patient characteristics are shown in **Table S1**. For each patient, 20-60 minutes of continuous NREM sleep was automatically selected^46^ and visually confirmed from each available night by a board-certified epileptologist. One exception was made for P52, for whom one night contained 17.9 minutes of continuous NREM sleep. The NREM segments were sampled during periods that were remote from seizure events (no seizures occurred within the 30 minutes preceding or following the sampled NREM segment). This resulted in a total of 336 nights of NREM data. On average, each patient had 6.11 ± 2.5 nights and 47.02 ± 12.5 continuous minutes of NREM, and 5.15 ± 2.7 thalamic subnuclei sampled. Of the 178.38 ± 43.93 bipolar channels acquired per patient (9811 channels in total), 8.45 ± 4.6 were located in the thalamus and 20.07 ± 15.2 were in the SOZ. SOZ channels ipsilateral to the thalamus were analyzed, and bilateral thalamic implantations were split into two separate observations, resulting in a total of 72 *patient-hemispheres*. Next, we grouped spindle detections into the six different groups of thalamic nuclei (ventralateral, VL, n=61; pulvinar-medial, PuM, n=22; pulvinar-others, PuO, n=24; anterior nucleus, ANT, n=17; centromedian, CM, n=39; and mediodorsal, MD, n=18) based on the Iglesias atlas^47^ with established FreeSurfer scripts^48^ (**Figure 1A**). This resulted in 2.96 ± 1.1 unique nuclear groups per patient-hemisphere. For subnucleus analysis, one patient was excluded since the localized thalamic contacts were not included in one of the six nuclear groups, resulting in 70 patient-hemispheres. The overall methodology to investigate long and short timescale interactions between the SOZ and thalamus is described in **Methods** and illustrated in **Figure 1**.

**Figure 1:**
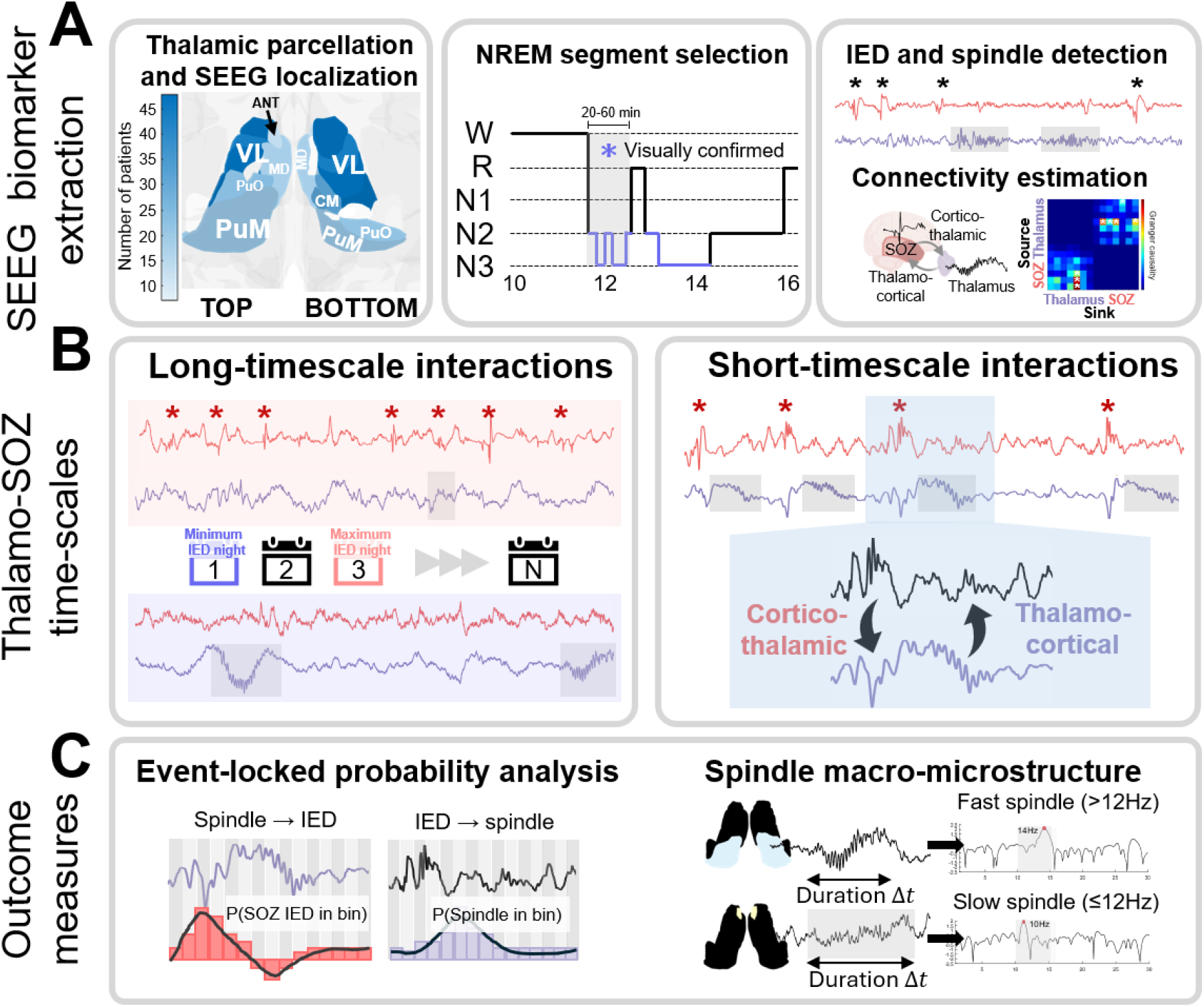
Methodology to investigate the effect of thalamocortical sleep dynamics on epileptic activity across short and long timescales. A. Pre-implantation T1 MRI was segmented using FreeSurfer^48^ to parcellate thalamic subnuclei using the Iglesias atlas^47^ and localize SEEG contacts. Thalamic contacts were assigned to thalamic subnuclei and grouped into six nuclear groups for analysis. Patient coverage across thalamic nuclear groups showed largest sampling in VL, CM, PuM and PuO nuclei. NREM sleep was automatically scored using SleepSEEG^46^, and candidate 20–60 min NREM segments were clipped and manually reviewed. If interruptions were identified during NREM, segments were re-clipped to ensure continuity. IEDs were detected^49^ in SOZ channels and sleep spindles were detected^50^ in thalamic channels. Connectivity between SOZ and thalamic bipolar channels was estimated using Granger causality^72^ to characterize directional thalamocortical and cortico-thalamic interactions. B. Effects of thalamocortical sleep dynamics on epileptic activity were analyzed across long and short timescales. Long-timescale analyses correlated thalamic spindle structure across contrasting nights of minimum and maximum IED rates in the SOZ. Short-timescale analyses quantified event-locked interactions between thalamic spindles and IEDs in the SOZ to assess the association of spindles with the changes in probability of IED occurrence. Red asterisks denote example detected IEDs, and gray shaded regions denote example detected spindles in the thalamus. C. Long-timescale analyses quantified changes in thalamic spindle rate and microstructure such as duration and central frequency. Short-timescale analyses quantified event-locked probabilities, such as the probability of SOZ-IEDs around thalamic spindle onset and the probability of thalamic spindle onset around SOZ-IED peaks. These measures were used to evaluate whether thalamic spindles are associated with subsequent changes in SOZ-IED probability, and whether SOZ-IEDs are associated with subsequent spindle facilitation. Abbreviations: SOZ, seizure onset zone; IED, interictal epileptiform discharge; SEEG, stereo-electroencephalography; NREM, non-rapid-eye movement; VL, ventrolateral; CM, centromedian; PuM, medial pulvinar; PuO, other pulvinar, ANT, anterior nucleus of the thalamus; MD, mediodorsal.

### Decreased IED activity is associated with increased spindle rates across long timescales

To investigate long-term interactions between the SOZ and the thalamus, we determined how spindle structure changed in nights of minimum IED rates compared to nights of maximum IED rates (**Figure 1B** right). We automatically detected IEDs^49^ in the SOZ and sleep spindles^50^ in the thalamus (**Figure 2A**). Across patient-hemispheres, the thalamic spindle rates in the night with minimum IEDs were substantially higher than in the night with maximum IEDs. The median percentage decrease in IED rates on the minimum IED night compared to maximum IED night was 51.9% (IQR, 39.9-67.3%), with an associated median 19.9% increase in spindle rates (IQR, -9.3.3-55.1%). We hypothesized that IED activity in the SOZ is associated with long-term connectivity changes to thalamic spindle structure (**Figure 1C** right). Therefore, IED rates were computed on (i) all SOZ channels; (ii) SOZ channels with strong connections *to* the thalamus; (iii) SOZ channels with strong connections *from* the thalamus; and (iv) thalamic channels (see **Methods**). Directed connectivity between the SOZ and thalamus was estimated using Granger causality^51^ (GC), which assesses whether activity in one brain region can predict future activity in another region (see **Methods**). Nights with the lowest IED rates in SOZ channels connected to the thalamus showed significantly increased thalamic spindle rates (Wilcoxon signed-rank test p<0.001, rank-biserial coefficient r=0.46, n=72; **Figure 2B**). Results were consistent for all SOZ channels (p=0.007, r=0.36, n=72) and SOZ channels connected from the thalamus (p=0.002, r=0.42, n=72). However, IED rates in the thalamus were not associated with changes to thalamic spindle rates (p=0.61, r=0.07, n=72; **Figure 2C**). Thus, subsequent analyses used the SOZ channels significantly connected to the thalamus, which is in line with nucleus-specific thalamocortical dynamics effective connectivity dependance^52^. Our nucleus-specific analysis revealed that the increase in spindle rates in nights with minimum IED rates was most pronounced in PuM (p=0.026, r=0.54, n=22) and VL nuclei (p=0.003, r=0.44, n=61) (**Figure 2D**). These results suggest that nights with reduced IED perturbations reflect a brain state associated with increased spindle activity and hence more stable sleep.

**Figure 2:**
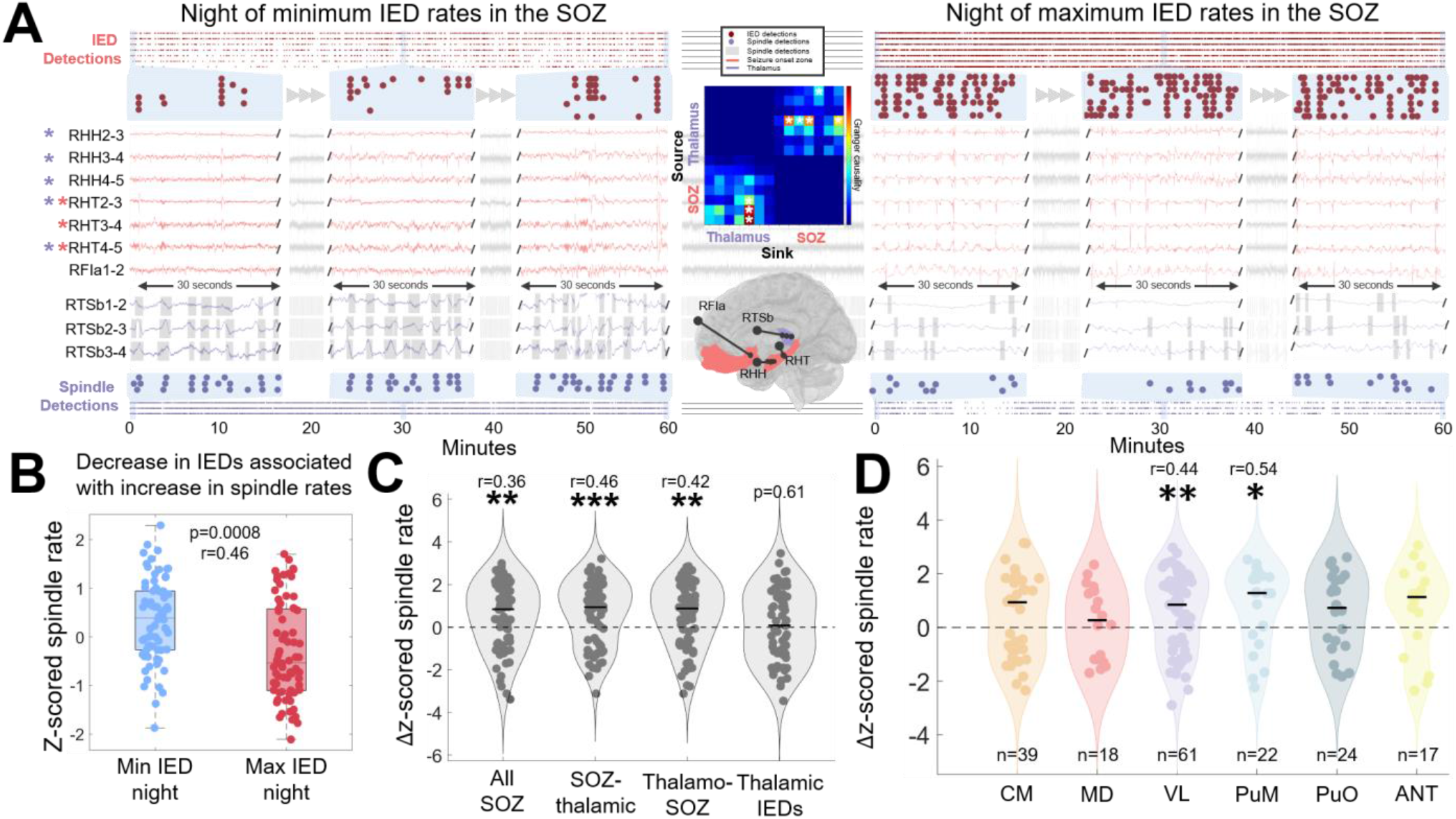
Thalamic spindle rates are higher on nights with minimum IED rates compared to nights with maximum IED rates. **A** Example of long-term interactions in patient P31 with SOZ located in the hippocampus and medial orbitofrontal gyrus. The thalamic SEEG was localized to the PuM, ANT and VL nuclei. In this illustration, only SEEG channels in the PuM were shown. IED detections are shown in dark red circles above the SEEG, and spindle detections are shown in purple dots below the thalamic SEEG. Gray shaded regions illustrate detected onset and offset of spindles in the thalamus. All SOZ channels are shown for illustration for two contrasting nights of IED activity. White asterisks indicate the GC connectivity that is above standard deviation of all positive values within the SOZ-thalamic or thalamo-SOZ connectivity. Red asterisks indicate the SOZ channels with SOZ-thalamic connectivity passing the threshold, and purple asterisks indicate SOZ channels with thalamo-SOZ connectivity that passes the threshold. Three 30-second windows of SEEG activity of the SOZ and thalamus are shown to illustrate changes not only in spindle rates but also in spindle microstructure across contrasting nights. **B** For each patient-hemisphere, spindle rates were z-scored across nights, and Δz-scored spindle rate was computed as the z-scored spindle rate on the minimum IED night minus the z-scored spindle rate on the maximum IED night within each patient-hemisphere. Positive values indicate increased spindle rates on minimum IED nights. Nights with lower SOZ IED burden showed significantly increased thalamic spindle rates. **C** The increase in thalamic spindle rate was observed when IED rates were computed using all SOZ channels, SOZ channels with directed connectivity to the thalamus and SOZ channels with directed connectivity from the thalamus, but not when IED rates were computed in the thalamus. **D** Nucleus-specific analysis showed that the increase in spindle rates was most pronounced in VL and PuM. Points indicate patient-hemispheres. Violin plots illustrate the distribution across patient-hemispheres. Horizontal bars indicate the group median. p-values were computed using paired comparisons between minimum and maximum IED nights. Abbreviations: SOZ, seizure onset zone; IED, interictal epileptiform discharge; VL, ventrolateral; PuM, medial pulvinar; PuO, other pulvinar; CM, centromedian; MD, medial dorsal; ANT, anterior nucleus of the thalamus.

To assess whether this relationship varied over the patient’s admission at the EMU, we fitted a linear mixed-effects model (LMM) across all recorded nights (see **Methods**). This sensitivity analysis showed that increased thalamic spindle rates were associated with lower IED rates (F = 8.2491, p = 0.0043), with no significant interactions between IED rate and day from admission (p = 0.15). Detailed LMM statistics are tabulated in **Table S2**. In the 31 patient-hemispheres where the minimum IED night occurred after the maximum IED night, 58.1% of patients had spindle rates higher on the minimum IED night (**Figure S2**). When the minimum IED night occurred earlier, the spindle rate increase occurred in 78% of patients. These results suggest that the changes in spindle rates are not only explained by the time of the patient’s stay in the EMU but rather fluctuate with IED rates.

### Decreased IED activity is associated with changes to spindle microstructure across long timescales

Next, we asked whether spindle microstructure was altered on nights of minimum IED rates compared to nights of maximum IED rates. We found that the spindle’s central frequency (p=0.005, r=0.38) and duration (p<0.001, r=0.46) significantly increased compared to nights of maximum IED rates (**Figure 3A**). Interestingly, the increase in the spindle’s central frequency was specific to the PuM (p=0.009, r=0.64, n=22; **Figure 3B**). In addition, the increase in spindle duration was observed in VL (p<0.001, r=0.55, n=61), PuM (p=0.004, r=0.77, n=22), PuO (p=0.023, r=0.57, n=24), and ANT (p=0.021, r=0.71, n=17; **Figure 3C**). Based on the spindle’s central frequency, we dichotomized spindles into fast spindles (>12 Hz) and slow spindles (≤12Hz). Fast spindle rates significantly increased on minimum IED nights (p<0.001, r=0.49; **Figure 3D**), particularly in the PuM (p=0.039, r=0.50, n=22; **Figure 3E-F**) and VL (p=0.0033, r=0.43, n=61; **Figure 3E-F**). In contrast, slow spindle rates did not significantly increase (p=0.083), and the change in fast spindle rates was significantly greater than the change in slow spindle rates (p=0.008, r=0.36).

**Figure 3.**
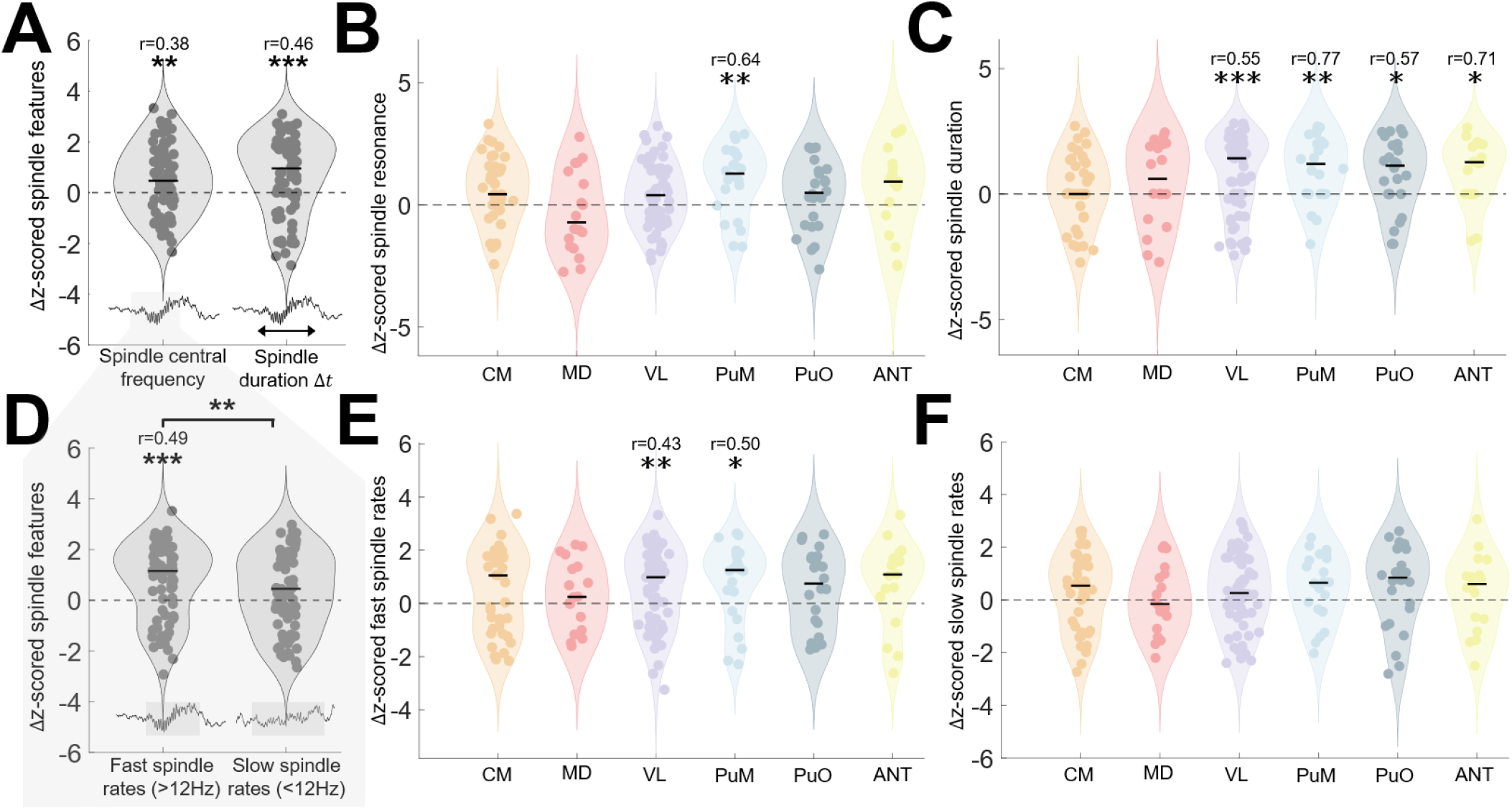
Decrease in IED rates are associated with nucleus-specific changes in spindle microstructure. **A** Changes in thalamic spindle central frequency and duration between minimum and maximum IED nights. Δ z-scored spindle features were computed as the z-scored value on the minimum IED night minus the z-scored value on the maximum IED night within each patient-hemisphere; therefore, positive values indicate increased values in spindle features on minimum IED nights. Both spindle central frequency and spindle duration increased on minimum IED nights. **B** The increase in spindle central frequency was observed across multiple thalamic nuclei, with the strongest effects in the PuM. **C** Spindle duration increased across several nuclei, with significant increases in the VL, PuM, PuO, and ANT. **D** Only fast spindle rates were larger on minimum IED nights. **E** Increase in fast spindle rates was most pronounced in VL and PuM, with **F** no changes found in slow spindle rates. Violin plots show distributions of paired within-patient-hemisphere differences; points indicate individual patient-hemispheres and black bars indicate group medians. Violin plots show distributions of spindle microstructures averaged over all nights, each point represents a patient-hemisphere. Asterisks indicate statistical significance. Abbreviations: SOZ, seizure onset zone; IED, interictal epileptiform discharge; VL, ventrolateral; PuM, medial pulvinar; PuO, other pulvinar; CM, centromedian; MD, medial dorsal; ANT; anterior nucleus of the thalamus.

### Individual spindles are followed by a decrease in post-spindle IEDs

Multi-night timescale analysis showed that decreased IED perturbations was associated with increased spindle rate, with higher central frequency and longer duration. Next, we asked how individual thalamic spindles influenced subsequent IEDs in the SOZ (i.e., hundred-milliseconds scale) (**Figure 1B** left). After time-locking spindle events to their onset, we estimated the probability an IED occurred around the time of spindle onset relative to baseline (**Figure 1C** left, see **Methods** for details). Baseline probability was computed as the average SOZ-IED probability -5 to -4 and 4 to 5 seconds away from thalamic spindle onset. We averaged all probability curves across channel pairs that have statistically significant non-zero GC in either direction between the SOZ and the thalamus. Following the spindle onset, we found that there was a significant decrease in IED probability (**Figure 4A**) that lasted around 500ms after spindle onset (FDR corrected p=0.002; r=0.41; at t=200ms from spindle onset). We also found a significant peak around 400ms before the spindle onset (FDR corrected p<0.001; r=0.86). This motivated our next analysis, which tested whether IED refractory periods could account for the acute decrease in IED probability (which we call ***IED suppression***) following spindle onset. This was estimated using time-locked intra-channel IEDs in the SOZ (see **Methods**). We found that 22 patient-hemispheres did not exhibit IED suppression following spindle onset. In the 50 patient-hemispheres with IED suppression, the decrease in IED probability following the onset of the spindle was significantly longer in duration than the upper bound of the empirical IED refractory period (p<0.001, r=0.67, n=50; **Figure 4A**, right panel).

**Figure 4:**
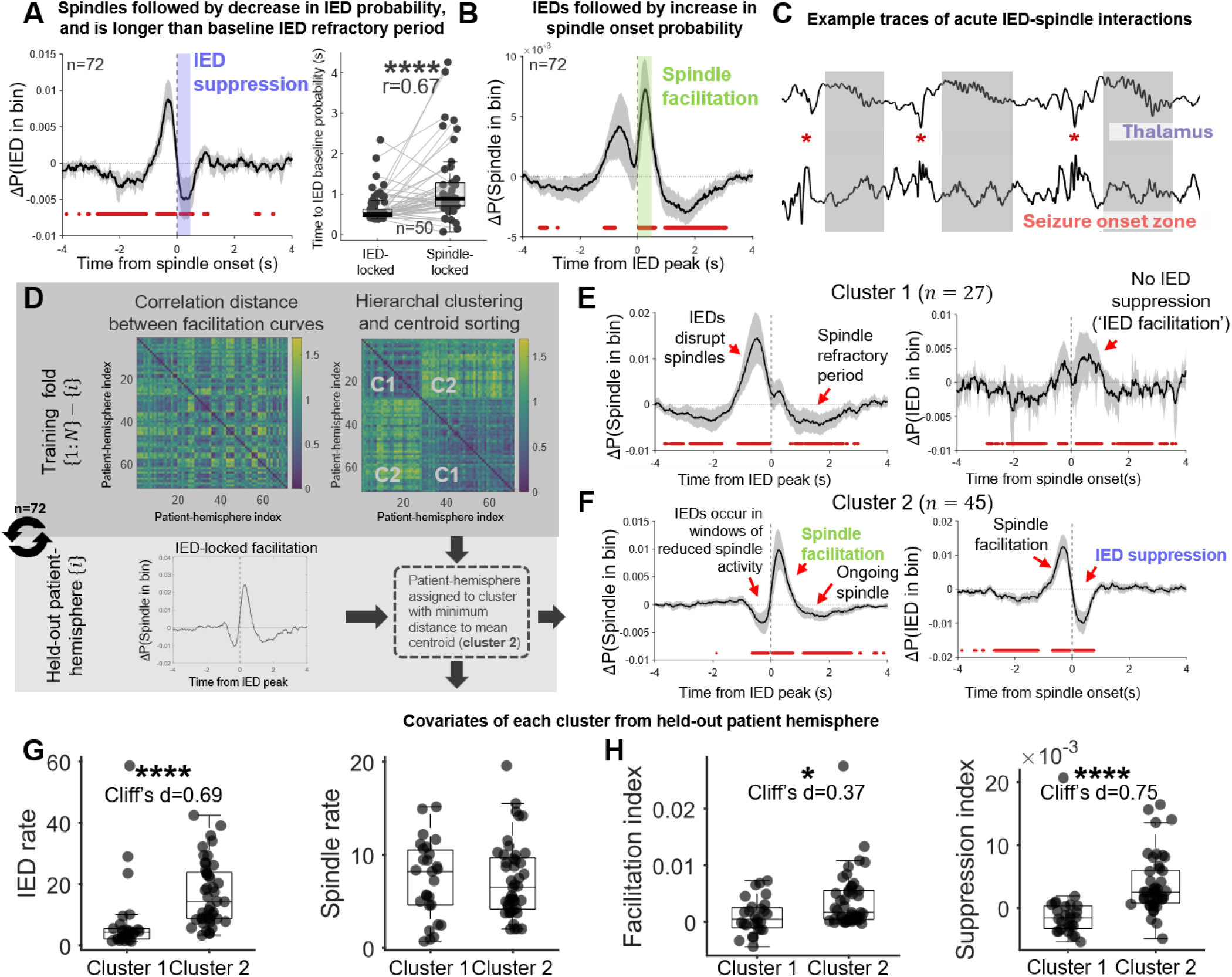
Spindle-locked IED suppression and IED-locked spindle facilitation are bidirectional processes. **A** Thalamic spindles were associated with a subsequent reduction in SOZ-IED probability, consistent with a short spindle-locked IED suppression period. Traces show the relative probability of IEDs around the spindle onset with respect to baseline probability [-5 -4s] and [4 5s]. Shaded regions indicate the 95% confidence interval. Red bars indicate time bins with significant deviation from baseline after Benjamini-Hochberg FDR correction, assessed by a two-tailed paired Wilcoxon signed-rank test. The inset compares the duration of spindle-locked IED suppression against the upper bound estimate of the IED refractory period using intra-channel SOZ-IED-locked IED probabilities. The results demonstrate that the duration of IED suppression from spindle-locked events is significantly longer than the IED refractory period in patients where spindle-locked IED suppression occurred (n=50). **B** IEDs in the SOZ were followed by an increase in thalamic spindle onset probability. **C** Representative SEEG traces showing acute IED-spindle interactions. Red asterisks indicate detected IEDs and gray shaded regions indicate detected thalamic spindles. **D** We employed a leave-one-patient-hemisphere-out approach to independently classify each left-out patient-hemisphere using the rest of the patients as training data. In each fold, pairwise correlation distances were computed between each patient-hemisphere IED-locked facilitation curve in the training set. We applied agglomerative hierarchical clustering using MATLAB packages **linkage** and **cluster**. The clusters were sorted by their mean centroids (t > 0) to ensure consistency in cluster IDs across each fold. The held-out patient-hemisphere was classified into either cluster based on the closest distance from its facilitation curve to the mean cluster centroids. The classification is done for each fold. **E** The increase in spindle probability prior to IED peak is only observed in cluster 1 and spindle-locked IED suppression did not occur in that cluster. **F** Cluster 2 exhibited the IED-locked spindle facilitation and spindle-locked IED suppression states. Interestingly, we see that the facilitation curve in cluster 2 preceding IED peaks show a significant decrease in spindle onset probability, likely explained by the fact that IEDs can only occur in windows where spindles do not occur, further supporting the bidirectional model of thalamo-SOZ suppression of IEDs. In addition, we see that spindle-locked IED suppression only occurs in patients which exhibited spindle facilitation as well, as shown in cluster 2. **G** We also compared covariates from each classified held-out patient across the two clusters. Patient-hemispheres in cluster 2 have significantly larger IED rates than patient-hemispheres in cluster 1 (Benjamini-Hochberg FDR correction). **H** As expected, we also found that the facilitation index and suppression indices are larger in cluster 2 compared to cluster 1 (Benjamini-Hochberg FDR correction). Together, these results suggest that increased sleep stability may exert a therapeutic effect on the SOZ. Abbreviations: SOZ, seizure onset zone; IED, interictal epileptiform discharge; FDR, false discovery rate

We then quantified the statistical significance of IED suppression following spindle onsets by fitting an LMM (see **Methods**) characterizing how IED probability varied with the spindle micro- and macro-structure (night-level median). The negative of the integral of the IED probability curve from 0 to 0.5 s after the spindle onset was calculated and defined as the suppression index (SI), such that positive values indicate large IED suppression following spindle onset. All channel pairs were considered irrespective of connectivity to investigate its interactions. Patient-hemisphere, night, and channel locations were considered random effects. Fixed-effects included the IED rates in the SOZ, spindle rates, spindle duration and central frequency, and whether the channel pair had statistically significant non-zero GC. We found that the predicted SI was statistically greater than zero (F = 13.22, ANOVA p<0.001) when all fixed-effect variables were fixed at their mean. This suggests that although the magnitude of IED suppression may vary with connectivity and microstructure, IED suppression is still a consistent effect and is not limited to specific conditions.

### IEDs are associated with an increase in post-IED spindles

Next, we aimed to identify interactions between IEDs in the SOZ and thalamic sleep spindles. Therefore, we time-locked SOZ-IED events at their peaks and estimated the probability a spindle onset occurs around the peak of an IED relative to baseline. We found that there was a significant increase in spindle probability following the peak of the IED (FDR corrected p<0.001, r=0.77, at t=325ms; **Figure 4B**), consistent with the increase in IED probability prior to spindle onset in **Figure 4A**. The integral of the spindle probability curve from 0 to 0.5 s after the peak of the IED was calculated and defined as the facilitation index (FI). We quantified the statistical significance of the increase in FI following the peak of the detected SOZ-IED by fitting the same LMM to the data as done previously. Indeed, the predicted FI was significantly greater than zero (F= 24.439, ANOVA p<0.001) when all fixed-effect variables were fixed at their mean. Illustrative sEEG signals examples show the interactions between thalamic spindle facilitation and subsequent suppression in SOZ-IED likelihood (**Figure 4C** and **Figures S3-5**).

### Spindle-locked IED suppression occurs exclusively in patients with IED-locked spindle facilitation

Interestingly, we saw a third phase in the facilitation curve preceding IED peaks, where there was a significant increase in spindle onset probability (**Figure 4B**). In addition, we found that IED suppression did not occur in 22 patient hemispheres. To analyze these findings further, we applied hierarchical clustering with a leave-one-patient-hemisphere-out approach (see **Methods**; **Figure 4D**). When splitting the dendrogram into two clusters, we found that cluster 1 represented patient-hemispheres that exhibited ‘spindle disruption’ (**Figure 4E**), while cluster 2 represented patient-hemispheres that exhibited ‘spindle facilitation’ (**Figure 4F**). Interestingly, we found that held-out patient-hemispheres that were grouped into cluster 2 (i.e., patient-hemispheres exhibiting IED-locked spindle facilitation) exhibited significant spindle-locked IED suppression (n=27), whereas patient-hemispheres in cluster 1 did not exhibit spindle-locked IED suppression (n=45). 18 of the 22 patient-hemispheres that were previously excluded (**Figure 4A** inset) were found in cluster 1. We also found that the patient-hemispheres in cluster 2 had statistically higher IED rates in the SOZ (p<0.0001, d=0.69, *n*_1,2_=27,45; median SOZ-IED rate = 14.37 min^-1^) compared to patient-hemispheres in cluster 1 (median SOZ-IED rate = 4.34 min^-1^; **Figure 4G**). Consistent with previous observations, SI (FDR corrected p<0.001; r=0.75) and FI (FDR corrected p=0.012; r=0.37) were both significantly higher in cluster 2 compared to cluster 1. Interestingly, we observed that the IED-locked spindle facilitation curve in cluster 2 had a significant decrease in spindle onset probability (**Figure 4F**), likely explained by the fact that IEDs can only occur in periods without detected spindles, further supporting the bidirectional model of thalamo-SOZ suppression of IEDs.

### Spindle microstructure shapes short-timescale cortico-thalamocortical interactions

Spindle microstructure was summarized as the median central frequency and median duration over all detected spindle events per night for each thalamic channel. Heatmaps of the FI and SI were generated by binning the central frequency and duration (see **Methods**). These heatmaps suggest a bimodal distribution in the FI and SI with respect to central frequency (**Figure 5A)**. Interestingly, spindle duration separated spindle facilitation from IED suppression, which was statistically significant across patient-hemispheres (p<0.001, r=0.90; (**Figure 5B**). To quantify this relationship, we identified the spindle-duration threshold that best separated IED suppression and spindle facilitation states, as illustrated in the heatmaps. A cutoff of 0.80 s was selected that optimized the AUC, and provided moderate discrimination (AUC = 0.73), where shorter spindle durations were associated with IED suppression and longer spindle durations were associated with spindle facilitation (**Figure 5C**). To quantify thalamocortical directional asymmetry, we then calculated the asymmetry index (ASI), defined as the normalized difference between the average thalamus→SOZ GC and SOZ→thalamus GC (see **Methods**). ASI ranges from -1 (thalamus→SOZ is stronger) to +1 (SOZ→thalamus is stronger). We found that stronger thalamic ASI (more negative) was associated with longer spindle duration (p<0.001, Spearman’s ρ=-0.42, n=72, **Figure 5D**), suggesting that thalamocortical asymmetry may reflect changes in the IED suppression/spindle facilitation state within the spindle-generating circuits.

**Figure 5:**
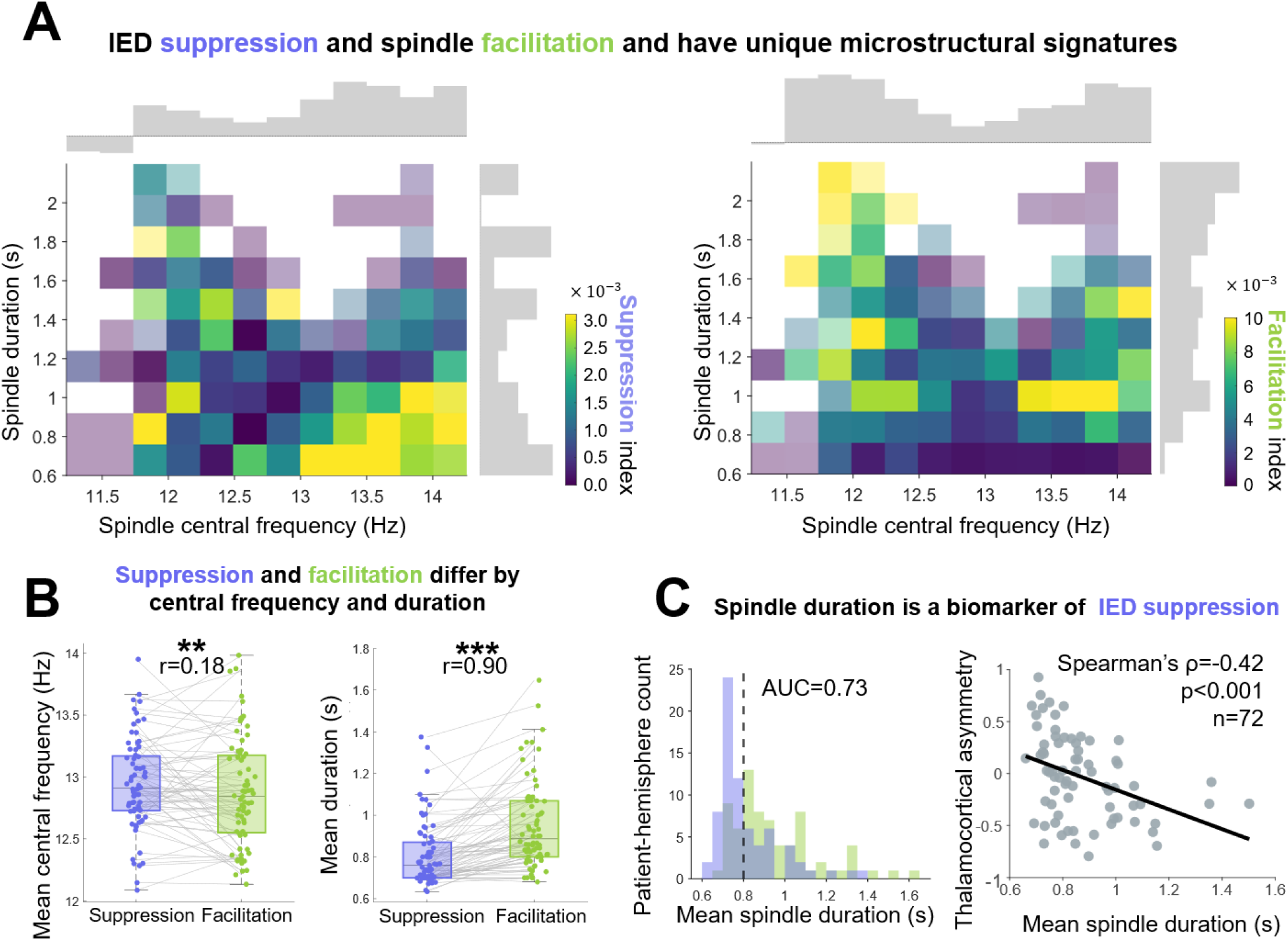
Short timescale bidirectional interactions between IEDs in the SOZ and spindles in the thalamus reveal distinct spindle microstructural states. A IED-locked spindle facilitation and IED suppression occupy distinct spindle microstructural states. Heatmaps show event-locked facilitation and suppression indices as a function of spindle central frequency and duration. Histograms illustrate the distribution of facilitation and suppression indices across spindle duration and central frequency. B Comparison of spindle microstructure between facilitation and suppression states. Spindle central frequency was significantly faster, and spindle duration was significantly shorter in the suppression regime. C Spindle duration separates suppression from facilitation states. A duration threshold classified suppression events with AUC=0.73, and shorter spindle durations were associated with stronger directional SOZ-thalamic asymmetry. Abbreviations: AUC, area under the curve; SOZ, seizure onset zone; IED, interictal epileptiform discharge

Consistently, the LMM revealed that the spindle central frequency was significantly associated with SI (F=9.5185, p<0.001), whereas spindle duration was not associated with SI (p=0.84). However, FI was associated with spindle central frequency band (F=11.923, p<0.001) and spindle duration (F=18.127, p<0.001). Although spindle duration did not correlate with changes in SI, it seemed to differentiate states of spindle facilitation and IED suppression. The fixed effects used in the LMMs are described in **Table S3**, and detailed results are tabulated in **Table S4**.

### Thalamic nuclei have distinct microstructural states that shape short-timescale cortico-thalamocortical interactions

Spindle microstructure across the six thalamic nuclear groups (ANT, CM, MD, PuM, PuO, VL) may contribute differently to IED suppression (**Figure 6A**) and spindle facilitation (**Figure 6B)**. We found that thalamic nuclei were significantly associated with FI (F=2.8221; ANOVA p=0.0149). While thalamic nucleus was not associated with SI (p=0.58), we found that its interaction with directed connectivity to the SOZ was significant: nuclei × thalamus→SOZ directed connectivity was significantly associated with SI (F=4.2055, ANOVA p<0.001), whereas nuclei×SOZ→thalamus directed connectivity was not statistically significant (p=0.41). Interestingly the inverse was true with FI: nuclei×thalamus→SOZ directed connectivity was not associated with FI (p=0.11), whereas nuclei×SOZ→thalamus directed connectivity was significant (F=3.8421, ANOVA p=0.0018). Together, these results suggest that directed connectivity does not strongly exert a uniform effect on IED suppression but is instead nucleus dependent and directional with respect to functional connectivity between the thalamus and SOZ.

**Figure 6:**
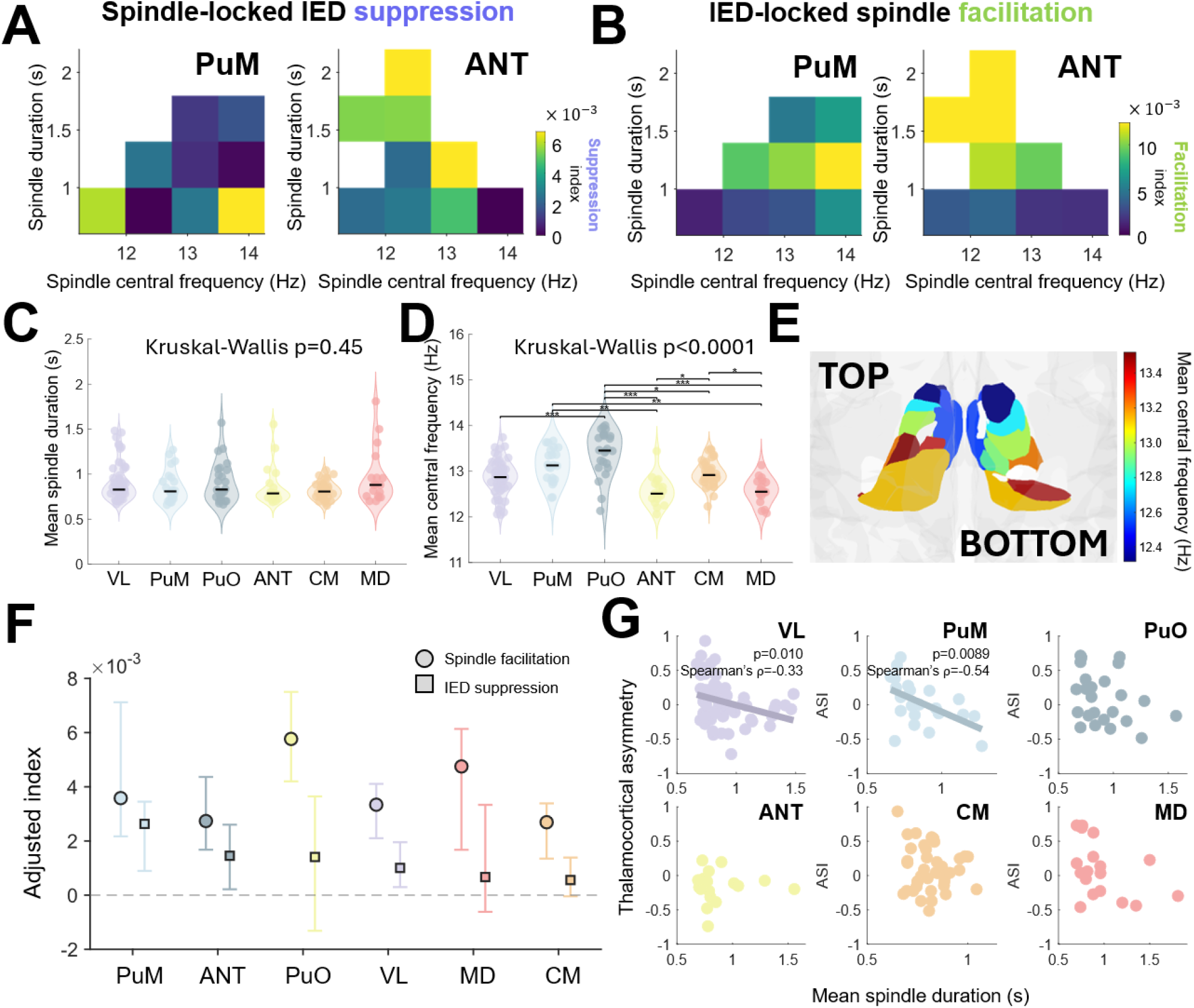
Short timescale IED–spindle interactions depend on nucleus and spindle microstructure. **A** spindle-locked IED suppression as a function of spindle central frequency and duration. Heatmaps illustrate the suppression index after thalamic spindle onset for PuM and ANT. **B** IED-locked thalamic spindle facilitation as a function of spindle central frequency and duration. Heatmaps illustrate the facilitation index for PuM and ANT. **C** Nucleus-specific distributions of spindle duration and **D** central frequency, demonstrating an **E** anteroposterior distribution of spindle central frequency. Violin plots show distributions of spindle microstructures averaged over all nights, each point represents a patient-hemisphere. **F** Adjusted suppression and facilitation indices were obtained from the linear mixed-effects model after fixing the fixed-effect variables at their means. Circles indicate IED-locked spindle facilitation and squares indicate spindle-locked IED suppression. Error bars denote the 95^th^ confidence interval estimated with bootstrapping (n=1000). **G** Relationship between spindle duration and thalamocortical asymmetry across nuclei. Shorter spindle durations were associated with stronger SOZ→thalamic directional asymmetry in VL and PuM nuclei, while other nuclei showed no significant association. Together, these results suggest that short timescale SOZ–thalamic interactions are nucleus-specific and depend on spindle microstructure, with spindle duration separating facilitation and suppression states. Abbreviations: SOZ, seizure onset zone; IED, interictal epileptiform discharge; VL, ventrolateral; PuM, medial pulvinar; PuO, other pulvinar; CM, centromedian; MD, medial dorsal; ANT; anterior nucleus of the thalamus.

Next, we compared spindle microstructure across thalamic nuclei. Spindle central frequency and duration were averaged over all nights for each patient-hemisphere. While we found that spindle duration did not vary across thalamic nuclei (Kruskal-Wallis p=0.45; **Figure 6C**), the spindle’s central frequency differed (Kruskal-Wallis p<0.0001, ε²=0.29; **Figure 6D**). Spindles recorded in PuM and PuO had a higher central frequency than spindles recorded in the ANT and MD, consistent with an antero-posterior organization of spindle frequency (**Figure 6E**) as known from their cortical distribution^53^.

Given that spindle central frequency varied across nuclei, we asked whether spindle microstructure could explain the variance in SI beyond nucleus-specific organization. By comparing LMM models with and without spindle duration and central frequency, we found that the LMM model that included spindle microstructure significantly improved model fit for SI (likelihood-ratio test = 28.784, p<0.001) and FI (likelihood-ratio test = 51.886, p<0.001). When fixing all fixed-effect variables at their mean, we found that the PuM had the largest adjusted SI, whereas ANT had the largest adjusted FI (**Figure 6F**), although they did not pass the statistical significance threshold. In addition, we found that greater thalamic ASI was associated with spindle duration in the PuM (p=0.0089, Spearman’s ρ=-0.54, n=22) and VL nuclei (p=0.010, Spearman’s ρ=-0.33, n=61), as shown in **Figure 6G**.

## DISCUSSION

This study demonstrates that spindles in the thalamus have bidirectional interactions with pathological disturbances and may help regulate network stability in epilepsy across multiple timescales. Across nights, we showed that spindle rates increased when IEDs decreased, with a stronger effect in SOZ channels connected to the thalamus. This inverse relationship did not systematically change over the course of patient’s admission and was associate with distinct spindle microstructural changes. Specifically, fast spindles (>12Hz) generated in the thalamus varied inversely with epileptic activity across multiple nights, potentially reflecting a protective inhibitory thalamocortical influence on the epileptic focus during NREM sleep that depends on spindle microstructure. This is consistent with our high-density scalp EEG study where we showed that fast spindle rates in early NREM had lower rates in the location of the epileptic focus compared to the contralateral side^54^.

We have shown for the first time that thalamic sleep spindles are associated with a transient decrease in IED probability that extends beyond the intrinsic refractory period of IEDs. This has significant implications as a potentially novel neuromodulation target to reduce epileptic activity while increasing sleep stability. While studies have already shown that IED perturbations can facilitate spindles in the cortex^37,42^, we have shown that IED suppression occurs in the context of spindle facilitation, and that spindles associated with IED suppression have distinct microstructures compared to spindles that are induced by IEDs (**Figure 5A**).

Interestingly, these interactions occurred in patients with prominent IED activity, irrespective of thalamic spindle activity. This suggests that sustained increases in IED activity may gradually remodel the thalamocortical network towards a state that is coupled with the SOZ, and might act as a potential covariate of thalamic neuromodulation efficacy. It may also explain why other studies have reported IED disruption in spindle structure^39^. Despite the increased IED rates in patient-hemispheres exhibiting spindle facilitation, spindle probability significantly decreased prior to the IED peak (**Figure 6**), which further supports the hypothesis that IEDs mostly occur in periods of reduced spindle activity. Moreover, these reciprocal interactions were associated with opposite directions of nucleus-specific thalamocortical connectivity. The probability of spindle-locked IED suppression was associated with nucleus-specific thalamus→SOZ connectivity, whereas the probability of IED-locked spindle facilitation was associated with nucleus-specific SOZ→thalamus connectivity. This directional asymmetry further highlights the importance of nucleus-specific effective networks for identifying thalamocortical neuromodulation targets. It has been shown that thalamocortical feedback can control spindle timing and duration^8^, consistent with our data demonstrating that directional thalamocortical asymmetry is correlated with spindle duration.

In long timescales, we demonstrated that spindle rates on nights of minimum IED rates increased the most in PuM. In short timescales, we showed, while not statistically significant, that the adjusted SI was highest in the PuM, whereas the adjusted FI was highest in the ANT. However, a larger sample size is needed to achieve the appropriate statistical power needed to identify unique involvement of different thalamic subnuclei in IED-spindle interactions. The study cohort consisted of patients with SOZs mostly located in temporal (49.1%) and parietal (5.5%) regions which was shown to form strong connections with the PuM nucleus^58–61^ and that response to stimulation depends on nucleus-specific effective connectivity to the SOZ^52,57^. This has direct implications for optimizing state-aware and patient-specific programming for thalamic neuromodulation in epilepsy.

These findings support a model in which IEDs and thalamic spindles interact through two opposing cortico-thalamocortical regimes^14^. IEDs generated in the SOZ perturb spindle-generating circuits which facilitate spindle generation in the thalamus, consistent with engagement of thalamic sleep circuitry, whereas spindles are followed by a period of IED suppression, which is significantly larger than the empirically derived IED refractory period. This further extends our current understanding of spindle-mediated protection against sleep disturbances^30,62^. We propose that the same thalamocortical mechanisms that maintain sleep stability may also contribute to the regulation of epileptic networks. Spindle generation has been shown to involve feedback oscillatory steady states that occur between the thalamic reticular nucleus and thalamocortical relay neurons^8,10,11^. The same circuit has also been implicated in promoting a seizure response^14^. Therefore, spindle activity may reflect a reticular-thalamic steady state that is shifted toward a more stable physiological regime, making the network less susceptible to pathological engagement. Therefore, we propose sleep spindles as a novel neuromodulation target (**Figure 7**) with the potential to reduce epileptic activity while also improving sleep stability^15,29^. This approach may be particularly valuable, given previous evidence that DBS of the ANT disrupts sleep in people with epilepsy^55–57^.

**Figure 7:**
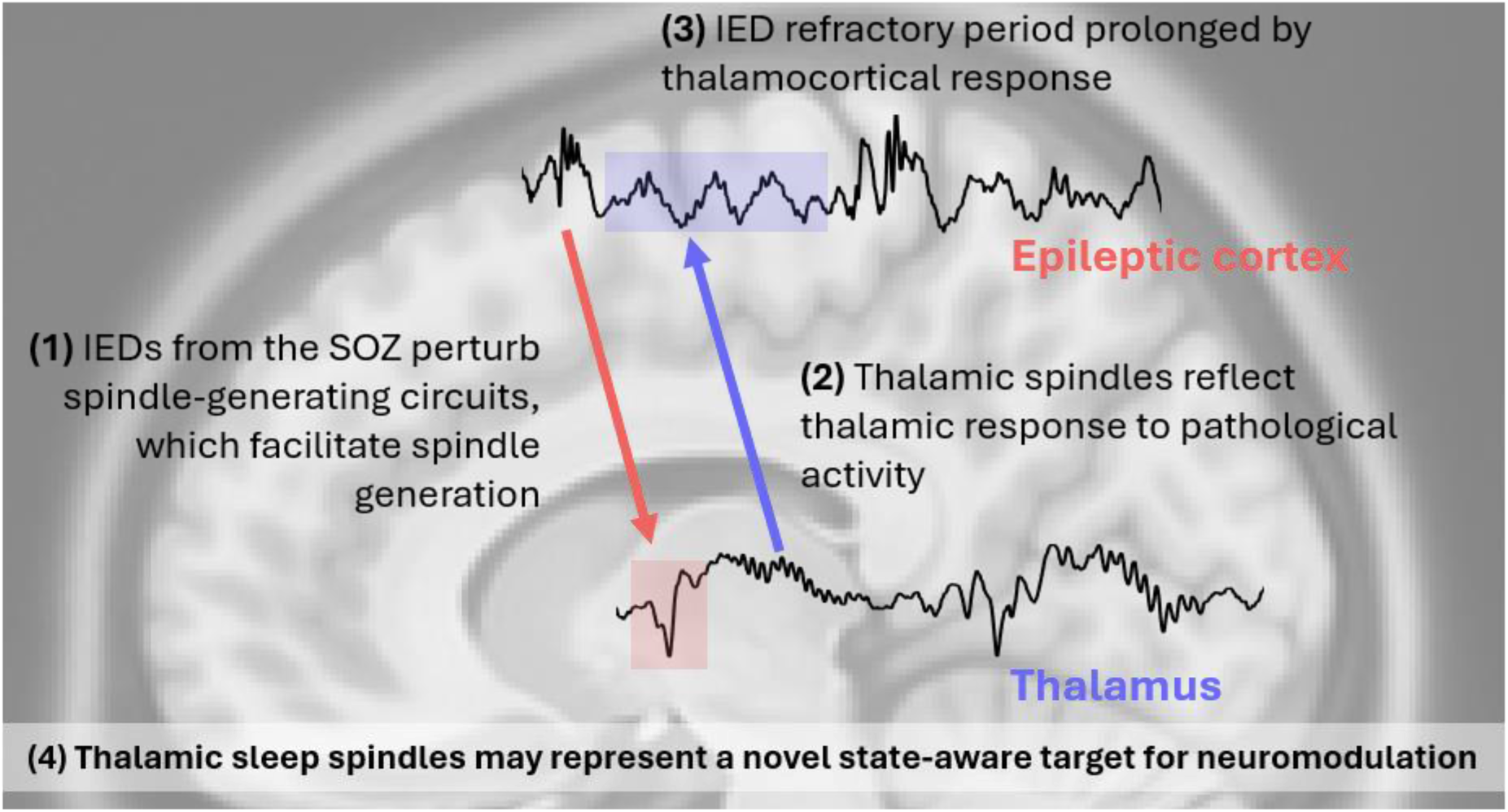
Theoretical model of bidirectional SOZ-thalamo-SOZ interaction. Our data suggests that IED perturbations may facilitate spindle generation, which in turn, may create a protective window against epileptic activity. Interestingly, spindle facilitation and IED suppression occurred only in patients with significantly higher IED rates. We also show that shorter spindles provide the protective window while IEDs facilitate the generation of longer spindles that occur in slow (<12Hz) and fast (>12Hz) spindle ranges. Unique spindle microstructural signatures involve different thalamic nuclei, suggesting that thalamic neuromodulation may require nucleus-specific optimization in future interventional studies aiming to induce the protective window for long-term reduction in epileptiform activity.

Our findings support the interictal suppression hypothesis which proposes that regions surrounding the epileptic focus may actively inhibit epileptic activity during interictal periods^63,64^. Our study demonstrates that IED activity from the epileptic focus may trigger a response in the thalamus, likely via thalamocortical feedback inhibition. This is consistent with IEDs inducing spindles in regions outside the seizure network^38^, and supports the idea that regions near the epileptic focus may be a subset of the overall distributed thalamocortical network that effectively suppress epileptic activity in the cortex via thalamocortical feedback inhibition^14^. Because thalamo-SOZ interactions varied across nights, this suppressive mechanism may fluctuate with brain state and contribute to variability in seizure expression^65^.

Although thalamic DBS and RNS are approved for the treatment of epilepsy^57,66^, they have not been proven effective for all patients. Our data provides a framework to help optimize neuromodulation outcomes by using patient-specific thalamocortical signatures to guide stimulation programming. The bimodal frequency structure observed across spindle facilitation and IED suppression may reflect how distinct spindle central frequencies differentially contribute to the IED-suppressive state. In addition, we observed a small but statistically significant increase in spindle central frequency on nights with the lowest IED rates. These results motivate future stimulation experiments to test whether entrainment at patient- and nucleus-specific spindle frequencies can selectively engage IED-suppressing thalamocortical networks. Spindle oscillations are sustained by repeated bursts of neuronal activity cycling through reticular, thalamocortical, and cortical populations^8^. The timing of these recurrent bursts contributes to the spindle central frequency and thus reflects the coordination of thalamocortical interactions. This coordination may be altered when reticular-thalamic circuits interact with epileptic cortex.

SEEG in epilepsy is the only way to study human thalamocortical interactions *in vivo*, yielding unique data that advances our knowledge of thalamocortical dynamics with implications for neuromodulation in human epilepsy. However, sEEG implantations are sparse with clinically biased spatial sampling of the epileptogenic network, resulting in measuring propagation regions rather than the true SOZ^67^. These limitations could contribute to variability in IED-spindle coupling across patients and hemispheres. We mitigated this issue by focusing the analysis on thalamo-SOZ channels with significant GC connectivity.

Since thalamo-SOZ interactions are the focus of this study, GC was only used to select channels that actively interact with each other. Finally, subgroup analyses across thalamic nuclei were limited by unequal sample sizes. The large number of thalamo-SOZ channel pairs and multinight recordings increased statistical power, although larger cohorts with greater representation of individual thalamic subnuclei will be needed to establish nucleus-specific effects.

Together, these findings extend our understanding of sleep spindles as more than just passive signatures of sleep. Our data demonstrate a bidirectional relationship between thalamic spindles and pathological activity, where pathological disturbances likely recruit spindle-generating circuits to regulate pathological excitability. These results establish a mechanistic foundation for state-aware thalamic neuromodulation approaches that harness sleep spindles to suppress pathological activity and maintain network stability.

## METHODS

### Participants

We retrospectively screened 58 consecutive patients who underwent phase 2 presurgical evaluation with sEEG at DUH (2023-2024) and MGH (2020-2024) with at least one sEEG electrode sampling the thalamus, and at least three nights of sEEG recordings available for analysis. Patients with non-focal SOZs were excluded. Of the 58 patients that were screened, three were excluded due to having less than three available nights (n=2) and a non-focal SOZ (n=1), resulting in a total of fifty-five patients included in this study (**Figure S1).**

### Stereo-EEG electrodes and localization

Patients at DUH who underwent sEEG presurgical investigation were implanted with PMT (PMT Corporation) and DIXI Medical (Besançon, France) electrodes. Patients at MGH were implanted with PMT (PMT Corporation) or Ad-tech (Racine WI, USA) depth electrodes. The DUH and MGH recordings were acquired using a Natus Quantum LTM amplifier (with Natus Neuroworks EEG software from Natus Medical, Inc.). DUH data was acquired with hardware filter cutoffs between 0.01 Hz and 812 Hz, and digitized at 2,048 Hz. MGH recordings were acquired with hardware filter cut offs between 0.01 and 340 Hz, and digitized at 1024 Hz. To harmonize the dataset across the different centers, DUH recordings were downsampled to 1024 Hz using the MATLAB function *resample*. All signals were re-referenced using a bipolar montage.

Following insertion of depth electrodes, patients underwent post-implantation CT imaging, which were then co-registered to pre-implantation T1 MRIs. Electrode reconstructions were manually performed on CURRY 8 for patients admitted to DUH and were performed in 3D Slicer for patients admitted to MGH. T1 MRIs were segmented and thalamic parcellations were computed using FreeSurfer scripts^47,48,68^. Three-dimensional coordinates of bipolar channels were constructed by taking the midpoint between adjacent contacts. Thalamic subnuclei were parcellated in FreeSurfer using the Iglesias atlas^47^. Each channel was assigned an anatomical region from the Iglesias atlas^47^ using an electrode labeling algorithm^68,69^. The Pulvinar nucleus was separated into two categories in accordance with deep-brain stimulation literature^60,66^. Nuclei were categorized into six different groups: VL, PuM, PuO, ANT, CM, and MD nuclei.

A board-certified epileptologist identified the SOZ for all patients, defined as channels showing the earliest unambiguous changes in the sEEG at seizure onset independent of the fast activity content and seizure-onset pattern^70^. We used the SOZ as a proxy of the epileptic tissue.

### Estimating connectivity between the thalamus and the SOZ

Our main hypothesis was that thalamic spindle activity interacts with SOZ-driven IED activity over different timescales. Consistent with literature on the importance of thalamo-SOZ hodology^52,58–60,71^, we estimated network connectivity between the thalamus and SOZ using GC^72^. GC is a statistical method that provides a data-driven measure to identify which SOZ channels propagate activity to or from the thalamus and has been used in previous studies to investigate thalamocortical connectivity in epilepsy^52,73,74^. It tests whether the past activity of a source *X*_1_ significantly improves the prediction of a sink *X*_2_. More formally, a connection *X*_1_ → *X*_2_is GC if the multi-variate autoregressive model using past values of *X*_1_(*t*) and *X*_2_(*t*) is significantly better at predicting *X*_2_(*t*) than using *X*_2_(*t*) alone:

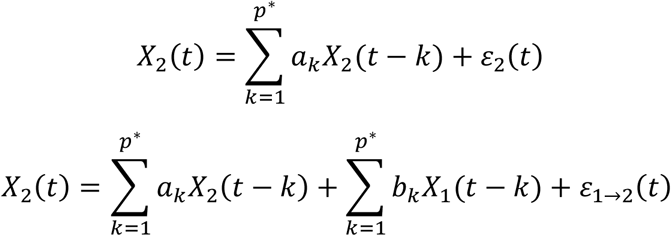

The GC value is the reduction in residual variance after including past values of *X*_1_:

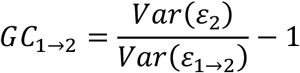

We used Brainstorm’s *bst_granger.m* implementation of GC^75^ in MATLAB. Signals were detrended, and the model order was optimized using Akaike’s final prediction error criterion with a maximum order of 10. Across all model-order estimates from 336 nights, the selected order had a median of 3 and an interquartile range of 3-4. Only 0.3% of estimates reached the upper bound of 10, suggesting that the specified search range was unlikely to constrain model-order selection. The first 10 minutes of NREM sleep were used to estimate within-night thalamo-SOZ connectivity. The signal was divided into five trials of two-minute segments each. This was computed for each night, and the median over all nights was used to quantify the patient-specific thalamo-SOZ connectivity for the long timescale analysis. The per-night patient-specific GC connectivity was used for the short timescale analysis.

We calculated a metric called the ASI, which quantifies the thalamocortical directional asymmetry, and is defined as follows:

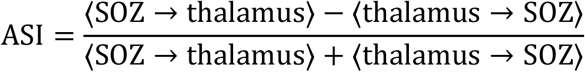

Where ASI ∈ {−1, +1} such that -1 represents stronger thalamus→SOZ connectivity, whereas +1 represents stronger SOZ→thalamus connectivity. ⟨SOZ → thalamus⟩ denotes the average GC connectivity from the SOZ to the thalamus, and ⟨thalamus → SOZ⟩ denotes the average GC connectivity from the thalamus to the SOZ.

We compared the asymmetry against spindle microstructure to explore how thalamocortical feedback may be reflected in the data.

Our hypothesis in this study is that functionally connected thalamo-SOZ networks exhibit inverse long-timescale dynamics. Therefore, for ***long timescale analysis*** we defined strong connections as those which deviate by more than one standard deviation of the distribution of non-zero connectivity values. More formally:

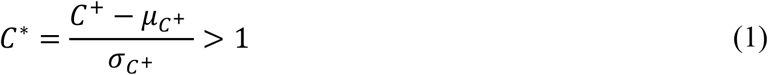

A threshold of one standard deviation was selected to include strong connections while preserving the total sample size, as some patients did not have connections above the threshold. For ***short timescale analysis***, we considered all non-zero connections from either direction in the analysis to investigate the short timescale bidirectional interactions. Although GC should not be interpreted as proof of causation, it serves as a useful metric to quantify directed connectivity. In this study, SOZ-thalamic GC was used to identify SOZ channels where past activity was predictive of thalamic activity, which is a data-driven way to study how SOZ-thalamic interactions relate to spindle macro- and micro-structure. In addition, thalamo-SOZ GC was used to identify thalamic channels were past activity predicted SOZ activity, which may reflect thalamic influence on the SOZ network during NREM sleep.

### Detection and characterization of IEDs and spindles

IEDs were detected in the SOZ, and sleep spindles were detected in the thalamus using previously validated detectors^49,50^. Detailed methodology for IED detection and post-processing is explained in our previous work^67^. IED rates were used to quantify the interictal epileptic burden for each night. Spindle duration and central frequency quantified spindle microstructure. Duration was computed as the time difference between detected offset and onset. The power spectrum was estimated from 1–30 Hz and flattened by subtracting the fitted 1/f background; the most prominent local maximum from 10–16 Hz defined the spindle’s central frequency.

### Sleep scoring and NREM segment selection

SleepSEEG was used to automatically sleep score overnight SEEG data^46^. Candidate NREM segments were chosen by selecting the earliest available continuous one-hour segment. If no continuous one-hour segment was available, the earliest segment with at least 20 continuous minutes of NREM sleep was selected. Candidate segments that were automatically selected were then visually reviewed for interruptions within the candidate NREM segment. In cases where wakefulness was found, the data was re-clipped to ensure continuity in NREM data. One night from P52 contained only 17.9 minutes of continuous NREM sleep and was retained as the exception.

### Investigating multi-scale interactions between SOZ and thalamus

The aim of this study was to investigate how thalamo-SOZ dynamics during NREM sleep are re-shaped across long and short timescales. This was done by (i) correlating IED burden with spindle micro- and macrostructure across multi-night data, and (ii) identifying spindle-locked and IED-locked changes to SOZ and thalamic activity.

#### Long timescale interactions

We compared spindle micro- and macrostructure across contrasting nights of minimum and maximum IED activity. IED activity was computed in SOZ channels in various conditions (i) all SOZ channels; (ii) SOZ channels that were GC connected to the whole thalamus; and (iii) SOZ channels that had GC connections from the thalamus. This was compared against IED activity detected within the thalamus to determine whether changes in spindle activity are specific to SOZ-IEDs. ‘Strong’ GC connections were determined using **Eq 1**.

To normalize the spindle features for group analysis, each feature was z-scored across all thalamus channels ipsilateral to the SOZ. This was done for the feature computed in the night of minimum IED rate and maximum IED rate for each condition.

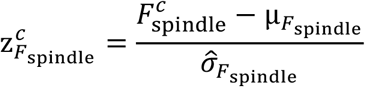

Where *c* ∈ {*i*_max_, *i*_min_} are nights of minimum and maximum IED activity, 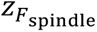 is the z-scored spindle feature *F*_spindle_, 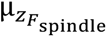 is the mean of *F*_spindle_ over all nights and all ipsilateral channels of interest, and 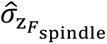 is the standard deviation of *F*_spindle_ over all nights and all channels of interest. To investigate how long-timescale changes to spindle macro- and micro-structure were influenced by different thalamic nuclei, spindle features from bipolar thalamic channels assigned to the same nuclear group were averaged within each patient-hemisphere and night.

#### Short timescale interactions

Next, we aimed to investigate cortico-thalamocortical feedback that may have contributed to changes in IED and spindle pathophysiology. We hypothesized that thalamo-SOZ influence is reflected by changes in IED probability following spindles, while SOZ-thalamic influence is reflected by changes in spindle probability following IEDs. Examples of these interactions in SEEG can be found in **Figures S3-4**.

We aimed to test this hypothesis by performing event-locked analysis. Spindles in the thalamus were time-locked to their onsets, whereas IEDs were time-locked to their peaks. IED probability was estimated by counting whether an IED peak occurred within a 250ms time bin with 95% overlap, between -5s to 5s relative to spindle onset. Therefore, each spindle had a Boolean vector of whether an IED occurred within each time bin and was time-locked to the spindle onset. This vector was averaged over all spindle events across the night within a thalamo-SOZ channel pair to estimate the conditional probability *P*(IED|spindle, *t*) for a given channel pair. This estimation was computed for each SOZ-thalamus pair across the complete NREM segment and then averaged across nights to obtain a single time-resolved probability for each patient-hemisphere. The same computation was done for IED-locked analysis, where spindle onsets were counted in 250ms time bins with 95% overlap between -5s to 5s relative to the peak of the IED. *For the overall IED probability curve, we only considered channel pairs with significant non-zero GC connections. For LMM modelling, all ipsilateral channel pairs were retained, and GC connectivity was modeled as a binary covariate. This was done to comprehensively investigate the importance of connectivity with respect to spindle-locked IED suppression and IED-locked spindle facilitation.* Each spindle-locked and IED-locked probability curve was subtracted by their baseline probability *P*_0_, where *P*_0_ is the baseline probability computed as the mean probability of spindle or IED occurring for each time bin between [-5 to -4] and [+4 to 5]s relative to event onset.

SI and FI were calculated as the integral of the relative probability curve from 0 to 0.5s post spindle onset and post IED peak, respectively. For SI, the negative of the integral was computed so that positive values reflect suppression. More formally:

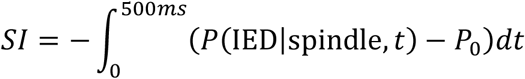

and

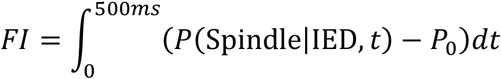

Next, we tested whether the duration of spindle-locked IED suppression was potentially related to the IED refractory period. We estimated two different endpoints (1) the time taken for spindle-locked IED suppression to return to baseline after spindle onset, and (2) the time taken for IED-locked IED probability to reach the nearest peak. For stable estimation of each endpoint, the probability curves were smoothed with a 250ms centered moving average to reduce noisy zero crossings. For the spindle-locked endpoint, suppression was required to begin within 250ms of spindle onset to allow tolerance for detector errors in marking the true spindle onset. The endpoint was then defined as the first time-point at which the spindle-locked IED probability curve returned to baseline. Regarding the IED-locked endpoint, we were unable to estimate the IED refractory period since IEDs were postprocessed to remove any IED event within 300ms around a given IED.

To rule out confounds due to this postprocessing step, our goal was to compare the duration of spindle-locked IED suppression to the time taken for the intra-channel IED-locked IED probability to return to baseline. Given the uncertainty introduced by the 300ms removal of IEDs, we could not directly estimate the time required for IED-locked IED probability to return to baseline. Therefore, an upper-bound estimate was computed as the time taken for the IED-locked IED probability to reach the first positive local maximum after the 300ms exclusion window. This comparison was performed for each patient-hemisphere. If the algorithm did not find any spindle-locked IED suppression, then the patient-hemisphere was excluded from this sub-analysis, as we aimed to test whether spindle-locked IED suppression was longer than the expected IED-to-IED refractory period.

### Statistical analysis

#### Long timescale interactions

The two-tailed Wilcoxon signed-rank test was used to assess statistical significance in spindle macro- and micro-structure across contrasting nights of minimum and maximum IED rate. Rank-biserial coefficient was used to quantify the paired effect across patient-hemispheres. Kruskal-Wallis test was used to assess statistical significance of spindle duration and central frequency across different thalamic nuclear groups. When the Kruskal-Wallis test was significant, pairwise post-hoc comparisons were performed and corrected using the Benjamini-Hochberg FDR procedure. A total of 15 comparisons were performed when comparing spindle central frequencies across the six groups of thalamic subnuclei.

We also aimed to support the analysis by fitting an LMM using the multinight data, and determine if IEDs within a patient-hemisphere (PatientHemiID) could explain the variance of spindle rates. This relationship was associated with the day from admission to investigate the potential effects of ASM tapering on the results. We fitted the multi-night (n=452 nights) data across all patient-hemispheres to the following linear mixed-effects model (LMM) structure:

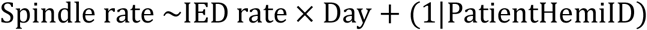

Where IED rates are night-level means computed over SOZ channels connected to the thalamus, and are zscored at the patient-hemisphere level across the nights. Day denotes the number of days after admission, which was z-scored across all entries, and PatientHemi represents the patient-hemisphere index which is considered the random effect.

#### Short timescale interactions

For the IED-locked spindle and spindle-locked IED probability curves, the two-tailed Wilcoxon signed-rank test was used followed by Benjamini-Hochberg FDR correction applied across all time-bins to identify time points that significantly deviated from baseline with a statistical threshold of 0.05. We then investigated the bidirectional relationship by clustering the IED-locked spindle facilitation curve in a leave-one-patient-hemisphere-out approach. We used an agglomerative approach, as it is a non-parametric method to group patient-hemispheres based on the temporal similarity of the facilitation curves. A correlation distance measure was used to characterize the similarity of each patient-hemisphere’s temporal profile. For each held-out patient-hemisphere *i*, the remaining 71 patient hemispheres were clustered into two groups using agglomerative hierarchical clustering with correlation distance and average linkage in MATLAB. We clustered the patient-hemispheres using the full time-support of the facilitation curve from -5 to 5 seconds. The clusters were then sorted based on the temporal mean (t > 0) to align them across each fold. The held-out patient-hemisphere was assigned to the cluster such that the training centroid had the smallest correlation distance from its facilitation curve. This resulted in obtaining a held-out patient-hemisphere facilitation and suppression curve for each fold. The corresponding held-out facilitation and suppression curves, as well as the electrophysiological covariates of the patient-hemispheres were then compared between the two clusters.

To investigate how thalamo-SOZ connectivity and spindle microstructure influenced spindle-locked IED suppression and IED-locked spindle facilitation, LMMs were used to test whether the effects were (1) different from baseline; (2) depended on spindle microstructure and thalamic nucleus; and (3) depended on directed connectivity between the thalamus and the SOZ. *Only for LMM modeling, all channel pairs were considered, and their directed connectivity was modeled as a binary covariate.* This was done to investigate how connectivity interacts with the other parameters with respect to SI and FI. Since the observations were repeated across nights (PatientHemiNightID), SOZ channels (SOZChannelID) and thalamus channels (ThalamusChannelID), they were modelled as random intercepts in the LMM. Inter-patient-hemisphere variability (PatientHemiID) was also accounted for by modelling it as a random intercept, as follows:

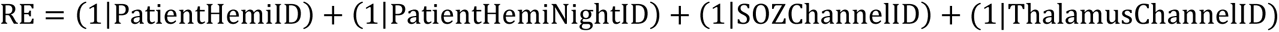

The following LMM model was fit to the data:

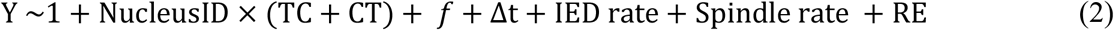

Where Y is either SI or FI. *f*, Δ*t*, TC and CT, represent the spindle central frequency (categorized into bins from 11 Hz to 15 Hz in steps of 1 Hz), spindle duration (continuous variable), TC (binary thalamo-SOZ connectivity), CT (binary SOZ-thalamic connectivity), and NucleusID (categorical variable representing six nuclear groups). Spindle rate (continuous variable) is the spindle rate of the thalamic channel, and IED rate (continuous variable) is the IED rate in the SOZ channel. Spindle central frequency was categorized into 1 Hz bins due to the bimodal nature of the SI and FI shown in the spindle microstructure heatmaps in **Figure 5A**. Adjacency matrices were binarized into 1 and 0, where 1 represents a non-zero statistically significant GC value between thalamus and SOZ pairs with p<0.05, and 0 represents a non-significant GC value. The median across all detected events was used to summarize the spindle microstructure for each thalamus channel, resulting in median central frequency and median duration for each thalamus channel of each night. All continuous variables were z-scored to ensure that predictors were on a similar scale.

To visualize the effects of various thalamic nuclei and spindle microstructure on IED suppression and spindle facilitation, we computed a heatmap of SI and FI against spindle central frequency and duration. A bin size of 12 and 10 was used to bin central frequencies and durations, respectively. Within each frequency-duration bin, SI or FI values were first averaged within each patient-hemisphere and then averaged across patient-hemispheres, to ensure that patient-hemispheres with more channel pairs or nights did not disproportionately influence the heatmap. For nucleus-specific analysis, a bin size of 4 was used to discretize spindle microstructure by FI and SI. For visualization purposes and to minimize visual effects of outliers, the color limits of each heatmap in **Figure 5A** and **Figure 6A-B** ranged from 0 to the 90^th^ percentile of the overall distribution.

To directly compare the spindle microstructure associated with spindle-locked IED suppression and IED-locked spindle facilitation, spindle microstructure was averaged within the upper 95th percentile of the SI and FI, respectively, for each patient-hemisphere. This resulted in one SI and one FI value for each patient-hemisphere. We statistically tested the difference in spindle duration and central frequency across FI and SI using a permutation test. This involved first identifying the SI and FI (averaged for each patient-hemisphere) within the top 5% of the total distribution. Before averaging the central frequency and duration values, they were shuffled and then averaged. For each patient-hemisphere, 10000 random permutations were performed. The two-sided permutation p-value was then computed as follows:

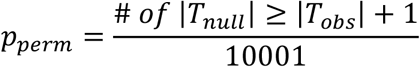

Where *T_null_* is the null distribution constructed for each patient-hemisphere, and *T_obs_* is the test statistic computed as the difference in mean central frequency and duration between IED suppression and spindle facilitation which was calculated previously within the upper 95th percentile of the FI and SI. The effect size was quantified using the rank-biserial coefficient between facilitation and suppression states.

## ACKNOWLEDGEMENTS

We wish to express our gratitude to the EEG technologists, clinical staff, and technicians at DUH and MGH for their support with clinical EEG and SEEG recordings. We also thank Mark Kramer and Catherine Chu for sharing the spindle detector code and providing guidance on its implementation with our dataset. This work was supported by the American Academy of Sleep Medicine Foundation (AASM Foundation) through a Bridge to Success Grant for Mid-Career/Senior Investigators (PI: Birgit Frauscher) as well as NIH funding NIA 5R01AG090302 (PI: Rina Zelmann).

## AUTHOR CONTRIBUTIONS

K.J. and B.F. conceived the study and experimental design. K.J. developed the analysis framework, performed the primary coding and statistical analyses, and prepared the manuscript and figures. K.J., A.H., H.Y., G.A., and K.S. contributed to dataset preparation, organization, and curation. K.J. and P.B.V. reviewed raw EEG recordings to ensure continuity of NREM sleep. T.A., P.R., L.W., M.K., C.C., P.K., M.R., D.S., P.H., S.S.C., P.S., D.C., R.Z., and B.F. contributed to data acquisition, clinical data resources, methodological input, and/or interpretation of results. R.Z. and B.F. provided study supervision. K.J. and B.F. drafted the manuscript. All authors critically revised the manuscript and approved the final version.

## CODE AVAILABILITY

All data analyses were performed using MATLAB R2023 and are available on our GitHub page (https://github.com/Lab-Frauscher/IED_Spindle_Interactions).

