## Supplementary Materials for "Sleep spindles stabilize human thalamocortical networks, inhibiting pathological disruptions"

### Supplementary Figures

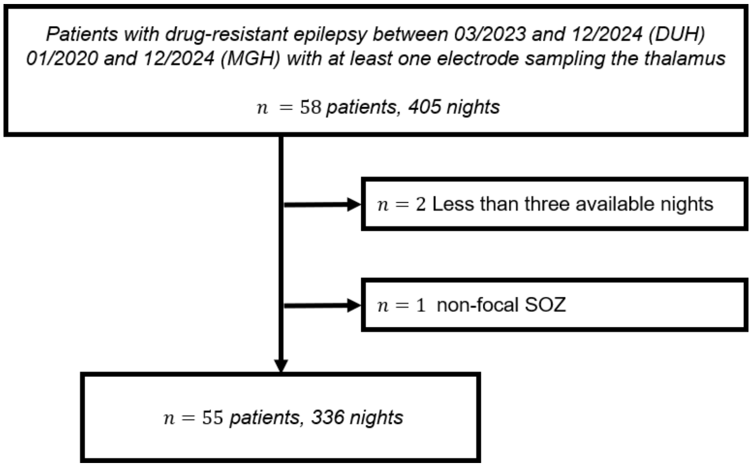


***Figure S1: Patient selection flowchart.*** *Of 58 screened patients, 55 were included based on the prespecified inclusion and exclusion criteria. Three patients were excluded due to the following reasons: less than three available nights with at least 20 minutes of continuous NREM data (n=2); and a non-focal SOZ (n=1)*

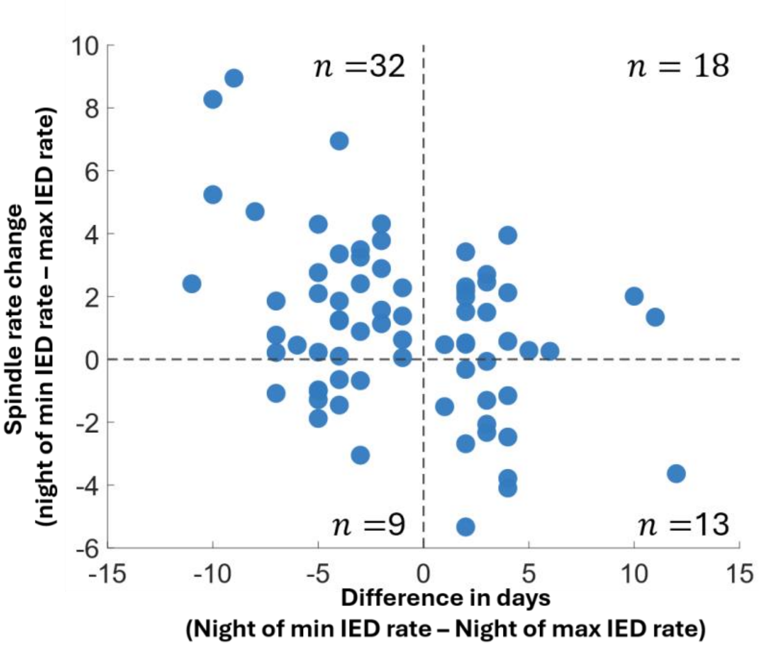


***Figure S2: Number of days between nights of minimum IED rate and maximum IED rate .*** *The number of days between the minimum and maximum IED nights was computed for each patient-hemisphere, and was plotted against the change in spindle rates. The increase in spindle rate on the night of minimum IED rates occurred in nights after the night of minimum IED rates for the majority (18/31) patients.*

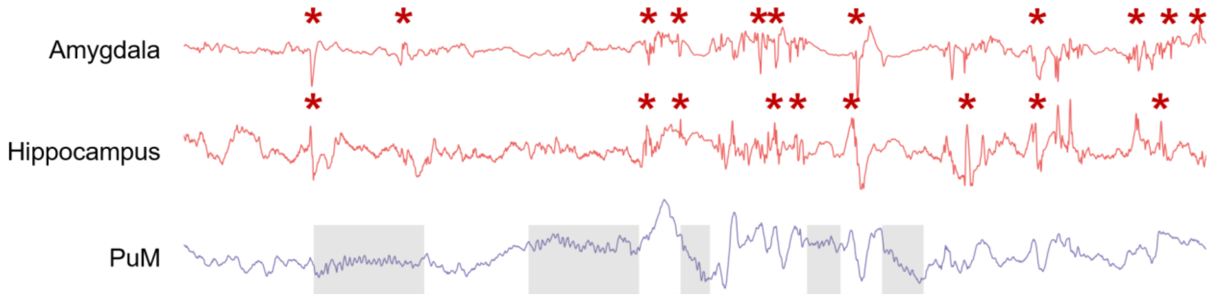


***Figure S3: Example SEEG of SOZ-thalamo-SOZ interactions during NREM sleep in patient P45.*** *SOZ is located in the mesial temporal lobe with the thalamus SEEG electrode targeting the medial pulvinar. Red asterisks denote IEDs in the SOZ, and shaded regions denote sleep spindles in the thalamus. Abbreviations: SOZ, seizure onset zone; IED, interictal epileptiform discharge; SEEG, stereo-electroencephalography; NREM, non-rapid-eye movement.*

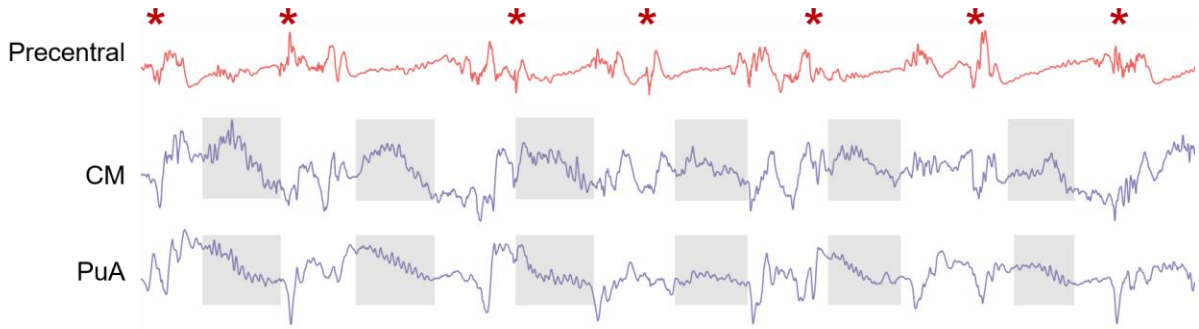


***Figure S4: Example SOZ-thalamo-SOZ interactions during NREM sleep in patient P17.*** *SOZ is located in the parietal lobe with thalamus SEEG electrodes targeting the centromedian and anterior pulvinar nuclei. Red asterisks denote IEDs in the SOZ, and shaded regions denote sleep spindles in the thalamus. Abbreviations: SOZ, seizure onset zone; IED, interictal epileptiform discharge; SEEG, stereotactic encephalography; NREM, non-rapid-eye movement.*

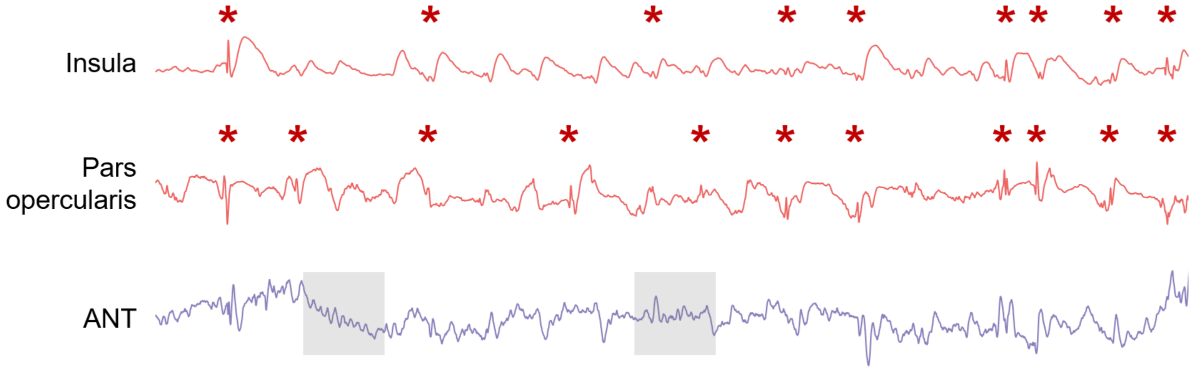


***Figure S5: Example SOZ-thalamo-SOZ interactions during NREM sleep in patient P40.*** *SOZ is located in the frontal lobe with the thalamus SEEG electrode targeting the anterior nucleus of the thalamus**.** Red asterisks denote IEDs in the SOZ, and shaded regions denote sleep spindles in the thalamus. Despite the persistent IED activity in the SOZ, note the unlikelihood of IEDs occurring during the spindle. Abbreviations: SOZ, seizure onset zone; IED, interictal epileptiform discharge; SEEG, stereotactic encephalography; NREM, non-rapid-eye movement.*

### Supplementary Tables

##### *Table S1: Patient demographics*

| Patient ID | Center | # Nights | Thalamic Nuclei | # Thalamus channels | # Electrodes | Minutes of NREM data | Electrode Names | SOZ Channels |
|---|---|---|---|---|---|---|---|---|
| P1 | DUH | 5 | CM, MD, VL | 9 | 18L | 59.7 | LFAI, LFAM, LFO, LFPI, LFPM, LIA, LIP, LPAI, LPAM, LPAS, LPMM, LTAI, LTAM, LTAS, LTMM, LTP, LTPM, LTPS | LTAI1, LTAI2, LTAI3, LTAI4, LTAM1, LTAM2, LTAM3, LTAM4, LTAM5, LTAM6, LTMM1, LTMM2, LTMM3, LTMM4, LTMM5, LTMM6, LTP1, LTP2, LTP3, LTP4, LTPM3, LTPM4, LTPM6 |
| P2 | DUH | 4 | PuO, PuM, VL | 6 | 9L+14R | 41 | LOPI, LPAI, LPI, LPPI, LTAM, LTMM, LTPI, LTPM, LTPS, ROAM, ROAS, ROPI, ROPM, ROPS, RPAI, RPI, RPPI, RPPM, RTAM, RTMM, RTPI, RTPM, RTPS | LOPI11, LOPI12, LTAM11, LTAM13, LTAM14, LTMM1, LTMM2, LTMM3, LTMM4, LTPI1, LTPI2, LTPI3, LTPI4, ROAM10, ROAM11, ROAM12, ROAM13, ROAM14, ROAM9, ROPI5, ROPI6, ROPI7, ROPI8, ROPM10, ROPM11, ROPM12, ROPM9, RPAI1, RPAI2, RTAM11, RTAM12, RTAM13, RTAM14, RTMM13, RTMM14, RTPI1, RTPI2, RTPI3, RTPI4 |
| P3 | DUH | 3 | PuO, VL | 7 | 15R | 46.83 | RAI, RFAM, RFAS, RFMM, RFMS, RFO, RFPI, RFPS, RPAI, RPAS, RPPI, RTAS, RTHA, RTHC, RTPS | RTAS3, RTAS4, RTAS5, RTAS6, RTPS3, RTPS4, RTPS5, RTPS6 |
| P4 | DUH | 6 | Unknown | 3 | 14L+5R | 50.92 | LFO, LIA, LPAM, LPPI, LPPM, LPPS, LTAI, LTAM, LTAS, LTMM, LTMS, LTP, LTPM, LTPS, RFO, RIA, RTAM, RTMM, RTPM | LTAI1, LTAI2, LTAM1, LTAM2, LTAM3, LTMM1, LTMM2, LTMM3, LTP1, LTP2, LTPM1, LTPM2, RTMM1, RTMM2, RTMM3, RTMM4, RTMM5, RTMM6, RTPM1, RTPM2, RTPM3, RTPM4 |
| P5 | DUH | 5 | PuO, PuM | 6 | 16L | 60 | LOAI, LOAM, LOAS, LOMI, LOMM, LOMS, LOPI, LOPM, LOPS, LPAI, LPPI, LTMM, LTOJ, LTPI, LTPM, LTPS | LOAI1, LOAI2, LOAI3, LOAI4, LOMI1, LOMI2, LOMI3, LOMI4, LOPI1, LOPI2, LOPI3, LOPI4, LTOJ1, LTOJ2, LTOJ3, LTOJ4 |
| P6 | DUH | 6 | PuO, PuM, VL | 4 | 3L+13R | 51 | LTAM, LTMM, LTPM, RFAI, RFAM, RFMM, RFO, RFPI, RTAI, RTAM, RTAS, RTMM, RTP, RTPI, RTPM, RTPS | RTAI2, RTAI3, RTAM1, RTAM2, RTMM1, RTMM2, RTPM3, RTPM4 |
| P7 | DUH | 15 | PuO, PuM, VL | 4 | 17L | 54.53 | LAI, LFAI, LFAM, LFO, LPI, LPPI, LTAI, LTAM, LTAS, LTLI, LTMI, LTMM, LTMS, LTP, LTPI, LTPM, LTPS | LTAI5, LTAI6, LTAS1, LTAS2, LTAS3, LTAS4, LTAS5, LTAS6, LTLI5, LTLI6, LTMI5, LTMI6, LTMS10, LTMS4, LTMS5, LTMS6, LTMS7, LTMS8, LTMS9, LTPS2, LTPS3 |
| P8 | DUH | 4 | CM, VL | 3 | 13L+2R | 49.04 | LFAI, LFAM, LFO, LIA, LTAI, LTAM, LTAS, LTMM, LTP, LTPI, LTPM, LTPS, LTTP, RTAM, RTMM | LTMM1, LTMM2, LTPM1, LTPM2 |
| P9 | DUH | 5 | CM, VL | 5 | 17L | 57.9 | LFAI, LFAM, LFAS, LFMS, LFOP, LFPM, LFPS, LPAI, LPAM, LPAS, LTAI, LTAM, LTAS, LTMM, LTP, LTPM, LTPS | LPAM1, LPAM2, LPAM3, LPAM4, LPAS1, LPAS2, LPAS3, LTAI1, LTAI2, LTAI3, LTAI4, LTAM1, LTAM2, LTAM3, LTAM4, LTMM1, LTMM2, LTMM3, LTMM4, LTMM5, LTMM6, LTP1, LTP2 |
| P10 | DUH | 11 | CM, Unknown, VL | 7 | 22L | 51.09 | LAIF, LAIP, LAIT, LAMP, LAMT, LANT, LASF, LASP, LAT, LCM, LINS, LMIF, LMIT, LMMT, LMSF, LMST, LOF, LPIF, LPIT, LPMT, LPSF, LPST | LAIP1, LAIP2, LAIP3, LAIP4, LAIP5, LAIP6, LAIP7, LAIP8, LAMP1, LAMP10, LAMP2, LAMP3, LAMP4, LAMP5, LAMP6, LAMP7, LAMP8, LAMP9, LAMT1, LAMT10, LAMT11, LAMT12, LAMT13, LAMT14, LAMT2, LAMT3, LAMT4, LAMT5, LAMT6, LAMT7, LAMT8, LAMT9, LAT1, LAT10, LAT11, LAT12, LAT13, LAT2, LAT3, LAT4, LAT5, LAT6, LAT7, LAT8, LAT9, LMIT1, LMIT10, LMIT11, LMIT12, LMIT2, LMIT3, LMIT4, LMIT5, LMIT6, LMIT7, LMIT8, LMIT9, LPIT1, LPIT2, LPIT3, LPIT4, LPIT5, LPIT6, LPIT7, LPIT8, LPMT1, LPMT2, LPMT3, LPMT4, LPMT5, LPMT6 |
| P11 | DUH | 3 | CM, VL | 4 | 18L | 60 | LFAI, LFAM, LFAS, LFMI, LFMM, LFMS, LFOP, LFP, LFPI, LFPM, LFPS, LTAM, LTAS, LTMI, LTMM, LTP, LTPM, LTPS | LTAM10, LTAM11, LTAM12, LTAM13, LTAM14, LTAM15, LTAM9, LTAS1, LTAS2, LTAS3, LTAS4, LTAS5, LTAS6, LTAS7, LTAS8, LTMI1, LTMI10, LTMI11, LTMI2, LTMI3, LTMI4, LTMI5, LTMI6, LTMI7, LTMI8, LTMI9, LTMM10, LTMM11, LTMM12, LTMM13, LTMM14, LTMM7, LTMM8, LTMM9, LTP1, LTP2, LTP3, LTP4, LTP5, LTP6, LTP7, LTP8, LTPM12, LTPM13, LTPM14, LTPM15, LTPS10, LTPS11, LTPS12, LTPS13, LTPS14, LTPS15, LTPS16, LTPS17, LTPS18, LTPS8, LTPS9 |
| P12 | DUH | 6 | PuO, PuM | 4 | 19L | 59.5 | A, B, C, LOAI, LOAM, LOAS, LOMI, LOMM, LOMS, LOPI, LOPM, LOPS, LPAI, LPI, LTMM, LTOJ, LTPI, LTPM, LTPS | LOAI3, LOAI4, LOAI5, LOAI6, LOAI7, LOAI8, LOMI5, LOMI6, LOMI7, LOMI8, LOPI10, LOPI5, LOPI6, LOPI7, LOPI8, LOPI9, LTPI10, LTPI3, LTPI4, LTPI5, LTPI6, LTPI7, LTPI8, LTPI9 |

| | | | | | | | | |
|---|---|---|---|---|---|---|---|---|
| P13 | DUH | 3 | PuM | 4 | 15L | 51.37 | LAI, LFAM, LFO, LFPM, LPAI, LPI, LTAM, LTAS, LTMM, LTMS, LTOS, LTP, LTPI, LTPM, LTPS | LPI11, LPI12, LPI13, LPI14, LPI15, LPI16, LTAM1, LTAM2, LTAM3, LTAM4, LTAM5, LTAM6, LTAM7, LTAM8, LTMM1, LTMM2, LTMM3, LTMM4, LTMM5, LTMM6, LTMM7, LTMM8, LTMS1, LTMS2, LTMS3, LTMS4, LTMS5, LTMS6, LTMS7, LTMS8, LTPI1, LTPI10, LTPI2, LTPI9, LTPM1, LTPM2, LTPM3, LTPM4, LTPM5, LTPM6 |
| P14 | DUH | 3 | CM, VL | 4 | 18R | 59.83 | RFAI, RFAM, RFAS, RFMI, RFMM, RFMS, RFOA, RFOB, RFPM, RIA, RTAI, RTAM, RTAS, RTMM, RTMS, RTP, RTPM, RTPS | RTAM1, RTAM2, RTAM3, RTAM4, RTAM5, RTAM6 |
| P15 | DUH | 8 | ANT, Unknown, VL | 6 | 16L+3R | 58.9 | LFAM, LFMM, LFO, LIA, LPMI, LPPI, LTAI, LTAM, LTAS, LTMM, LTMS, LTP, LTPI, LTPM, LTPO, LTPS, RTAM, RTMM, RTPM | LTAM1, LTAM2, LTAM3, LTAM4, LTMM1, LTMM2, LTMM3, LTP1, LTP2, LTP3, LTP4, LTP5, LTP6, LTP7, LTP8 |
| P16 | DUH | 6 | PuO, PuM | 4 | 17L | 60 | LOPI, LOPM, LPI, LPMM, LPPI, LTAI, LTAM, LTAS, LTLI, LTMI, LTMM, LTMS, LTPI, LTPM, LTPO, LTPS, X | LOPI7, LOPI8, LOPI9, LTLI5, LTLI6, LTPI7, LTPI8 |
| P17 | MGH | 6 | CM, PuO, VL | 11 | 5L+5R | 59.83 | LCM, LMa, LMb, LMc, LMd, RCM, RMa, RMb, RMc, RMd | LMa1, LMa2, LMa3, LMa4, LMc10, LMc7, LMc8, LMc9, RMb1, RMb2, RMb3, RMb4, RMb5, RMc10, RMc6, RMc7, RMc8, RMc9, RMd10, RMd6, RMd7, RMd8, RMd9 |
| P18 | MGH | 7 | CM, MD, VL | 10 | 4L+12R | 56.69 | LCM, LH, LPI, LPS, RCM, RFIa, RFIb, RFM, RFS, RH, RPIa, RPIb, RPSa, RPSb, RTM, RTS | LH1, LH2, LH3, LH4, LH5, RFIa5, RFIa6, RFIa7 |
| P19 | MGH | 3 | CM, MD, VL | 6 | 4L+9R | 59.33 | LFIa, LFIb, LFM, LPI, RFIa, RFIb, RFM, RPIa, RPIb, RPSa, RPSb, RPSc, RPSd | RPSc6, RPSc7, RPSc8 |
| P20 | MGH | 7 | CM, PuO, PuM, VL | 9 | 7L+7R | 59.57 | LF, LO, LPI, LPO, LPSa, LPSb, LTO, RF, RO, RPI, RPO, RPSa, RPSb, RTO | LPSa1, LPSa2, LPSa3, LPSa4, LPSa5, LPSa6, LTO10, LTO11, LTO12, LTO7, LTO8, LTO9, RF1, RF2, RF3, RF4, RPO1, RPO2, RPO3, RPO4, RPSa10, RPSa11, RPSa12, RPSa8, RPSa9, RPSb1, RPSb2, RPSb3, RPSb4, RPSb5, RPSb6, RPSb7, RPSb8 |
| P21 | MGH | 9 | CM, VL | 4 | 8L+8R | 59.33 | LCM, LFM, LFSa, LFSb, LFSc, LPI, LPSa, LPSb, RCM, RFM, RFSa, RFSb, RFSc, RPI, RPSa, RPSb | LCM1, LCM2, LCM3, LCM4, LCM5, LFM10, LFM11, LFM12, LFM13, LFM14, LFM7, LFM8, LFM9, LFSa1, LFSa2, LFSb2, LFSb3, LFSb4, LFSb5, RCM1, RCM2, RCM3, RFM10, RFM11, RFM12, RFM13, RFM14, RFSa1, RFSa2, RFSa3, RFSa4, RFSb1, RFSb2, RFSb3, RFSb4 |
| P22 | MGH | 6 | MD, Unknown | 5 | 7L+7R | 31 | LCM, LFSa, LFSb, LFSc, LH, LPI, LTM, RCM, RFSa, RFSb, RFSc, RH, RPI, RTM | LFSb1, LFSb2, LFSb3, LFSc10, LFSc11, LFSc12, LFSc4, LFSc5, LH2, LH3, LH4, LPI1, LPI2, RFSa10, RFSa11, RFSa12, RFSa6, RFSa7, RFSa8, RFSa9, RFSb10, RFSb11, RFSb12, RFSc10, RFSc11, RFSc12, RFSc8, RFSc9, RH3, RH4 |
| P23 | MGH | 5 | CM, MD, PuO, VL | 8 | 13R | 59.4 | RAMY, RFI, RFM, RFSa, RFSb, RHH, RHT, RPI, RPS, RTB, RTP, RTSa, RTSb | RAMY1, RAMY2, RAMY3, RAMY4, RAMY5, RHH1, RHH2, RHH3, RHH4, RHH5, RHH6, RHT1, RHT2, RHT3, RHT4 |
| P24 | MGH | 8 | CM, PuO, VL | 10 | 7L+9R | 31 | LAMY, LFI, LHH, LHT, LPIa, LPIb, LTO, RFI, RH, ROI, ROS, RPIa, RPIb, RPS, RTB, RTO | ROI10, ROI11, ROI7, ROI8, ROI9, RTB1, RTB2, RTB3, RTB4 |
| P25 | MGH | 6 | CM, VL | 5 | 10L | 31 | LCM, LFIa, LFIb, LOI, LOS, LPIa, LPIb, LPIc, LPS, LTI | LTI1, LTI10, LTI11, LTI12, LTI2, LTI3, LTI4, LTI9 |
| P26 | MGH | 12 | ANT, PuO, PuM, VL | 9 | 7L+1R | 31 | LAMY, LHH, LHT, LT, LTB, LTO, LTS, RTB | LHH1, LHH2, LHH3, LHH4, LHT1, LHT2, LHT3, LHT4, RTB5, RTB6, RTB7 |
| P27 | MGH | 9 | ANT, MD, PuO, PuM, VL | 13 | 8L+8R | 30.89 | LAMY, LFIa, LFIb, LH, LOS, LPI, LTB, LTS, RFIa, RFIb, RHT, ROS, RPIa, RPIb, RTB, RTS | LH1, LH2, LH3, RFIa10, RFIa11, RFIa12, RFIa13, RFIa14, ROS14, ROS15, ROS16, RPIb10, RPIb11, RPIb12, RPIb13, RPIb14, RTS15, RTS16 |
| P28 | MGH | 5 | ANT, VL | 7 | 8L+8R | 31 | LAMY, LFIa, LFIb, LFM, LHH, LHT, LTB, LTP, RAMY, RFIa, RFIb, RFM, RHH, RHT, RTB, RTP | LHH1, LHH2, LHH3, LHH4, LHH5, LHH6, LHT1, LHT2, LHT3, LHT4, LHT5, LHT6, LHT7 |
| P29 | MGH | 5 | ANT, CM, MD, PuO, PuM, VL | 20 | 7L+7R | 30.3 | LFIa, LH, LPIa, LPIb, LPIc, LPSa, LTS, RFIa, RFIb, RFIc, RFMa, RFMb, RPI, RTS | LFIa1, LH1, LH2, LH3, LH4, LH5, LH6, LH7, LPIa1, LPIa2, LPIa3, LPIa4, LPIb1, LPIb2, LPIb3, LPIb4, LPIb5, LPSa10, LPSa11, LPSa8, LPSa9, LTS10, LTS11, LTS12, LTS13, LTS14, LTS8, LTS9 |
| P30 | MGH | 5 | CM, VL | 4 | 11L+3R | 47.83 | LAMY, LFIa, LFIb, LFMa, LFMb, LFSa, LFSb, LHB, LHT, LTSa, LTSb, RAMY, RHB, RHT | LFIb1, LFIb2, LFIb3, LFMa1, LFMa2, LFMa3, LFMa4, LFMa5, LFSa1, LFSa2, LFSa3, LFSa4, LFSa5, LFSb2, LFSb3, LFSb4, LFSb5, LFSb6, LHT11, LHT12, LHT13, LHT14 |

| | | | | | | | | |
|---|---|---|---|---|---|---|---|---|
| P31 | MGH | 7 | ANT, MD, PuO, PuM, VL | 14 | 7L+8R | 59.42 | LFIa, LFIb, LHH, LHT, LTB, LTSa, LTSb, RAMY, RFIa, RFIb, RHH, RHT, RTB, RTSa, RTSb | LFIa5, LFIa6, LFIa7, RFIa1, RFIa2, RHH2, RHH3, RHH4, RHH5, RHT2, RHT3, RHT4, RHT5 |
| P32 | MGH | 6 | CM, Unknown, VL | 10 | 7L+7R | 50.56 | LCM, LH, LOI, LOS, LP, LTB, LTO, RCM, RH, RPI, RPS, RTB, RTO, RTS | LTB1, LTB2, LTB3, LTB4, LTB5, LTO1, LTO2, LTO7, LTO8, RTB1, RTB10, RTB2, RTB3, RTB4, RTB5, RTB6, RTB7, RTB8, RTB9 |
| P33 | MGH | 3 | CM, MD, VL | 9 | 8L+8R | 59.33 | LFIa, LFIb, LH, LPIa, LPIb, LPS, LTSa, LTSb, RFIa, RFIb, RH, RPIa, RPIb, RPS, RTSa, RTSb | LFIb1, LFIb2, LFIb3, LFIb4, LPIa1, LPIa2, LPIa3, LPIa4, LPIa5, LPIb12, LPIb13, LPIb14, LPIb15, LPIb16, LPIb6, LPIb7, LPIb8, LPIb9, RFIb1, RFIb2, RFIb3, RFIb4, RH1, RH2, RH3, RH4, RPIa1, RPIa10, RPIa11, RPIa12, RPIa2, RPIa3, RPIa4, RPIa5 |
| P34 | MGH | 7 | CM, VL | 9 | 3L+3R | 54.40 | LFI, LFM, LFS, RFI, RFM, RFS | LFM12, LFM13, LFM14, LFM15, LFM16, LFS12, LFS13, LFS14, LFS15, LFS16, LFS17, LFS18, RFM10, RFM11, RFM12, RFM13, RFM14, RFS10, RFS6, RFS7, RFS8, RFS9 |
| P35 | MGH | 7 | CM, PuO, VL | 9 | 5L+11R | 54.57 | LAMY, LFMa, LFMb, LHH, LHT, RAMY, RFI, RFM, RH, RPIa, RPIb, RPSa, RPSb, RTB, RTSa, RTSb | LHT1, LHT2, LHT3, LHT4, LHT5, RAMY3, RAMY4, RAMY5, RH1, RH2, RH3, RH4, RH5, RH6, RTSa1, RTSa2, RTSa3 |
| P36 | MGH | 7 | ANT, CM, MD, VL | 16 | 10L+4R | 59.14 | LAMY, LFIa, LFIb, LFIc, LH, LPIa, LPIb, LTB, LTSa, LTSb, RFIa, RFIb, RH, RPI | LFIa2, LFIa3 |
| P37 | MGH | 7 | ANT, VL | 6 | 8L+8R | 51.14 | LAMY, LFIa, LFIb, LH, LPI, LTB, LTSa, LTSb, RAMY, RFIa, RFIb, RH, RPI, RTB, RTSa, RTSb | LH2, LH3, LH4, LTB2, LTB3, LTB4, RH2, RH3, RH4, RH5 |
| P38 | MGH | 5 | CM, MD, VL | 4 | 10L | 30.3 | LAMY, LFIa, LFIb, LHH, LHT, LPIa, LPIb, LTB, LTSa, LTSb | LAMY1, LAMY2, LAMY3, LHH1, LHH2, LHH3, LHH4, LHT1, LHT2, LHT3 |
| P39 | MGH | 3 | CM, MD, VL | 6 | 11R | 31 | RAMY, RFIa, RFIb, RHH, RHT, RPIa, RPIb, RTB, RTSa, RTSb, RTSc | RFIa1, RFIa2, RFIa3, RFIa4, RFIa5, RFIa6 |
| P40 | MGH | 5 | MD, VL | 5 | 11L | 59 | LAMY, LFIa, LFIb, LFIc, LFId, LFM, LHH, LHT, LTB, LTSa, LTSb | LFIc10, LFIc11, LFIc12, LFIc13, LFIc14, LFIc15, LFIc16, LFId10, LFId11, LFId12, LFId13, LFId14 |
| P41 | MGH | 9 | ANT, CM, MD, VL | 20 | 12L+6R | 59.33 | LAMY, LAMYmicro, LFIa, LFIamicro, LFIb, LFMa, LFMb, LFS, LHH, LHT, LTB, LTS, RFIa, RFIb, RFMa, RFMb, RFS, RH | LFIa5, LFIa6, LFIa7, LFIa8, LFMa10, LFMa11, LFMa12, LFMa13, LFMa9, LFMb10, LFMb11, LFMb12, LFMb13, LFMb14, LFMb8, LFMb9, LFS10, LFS11, LFS12, LFS8, LFS9, LHH1, LHH2, LHH3, LHH4, LHH5, LHH6, LHT13, LHT14, LHT15, LHT16, LTB12, LTB13, LTB14, LTB15, LTB16, RFIb13, RFIb14, RFIb15, RFIb16, RFMa10, RFMa11, RFMa8, RFMa9, RFMb10, RFMb11, RFMb12, RFMb13, RFMb14, RFMb15, RFMb8, RFMb9, RFS10, RFS11, RFS12, RFS6, RFS7, RFS8, RFS9 |
| P42 | MGH | 4 | CM, PuO, PuM, VL | 20 | 7L+8R | 57.58 | LFI, LFM, LFSa, LFSb, LPI, LPS, LTB, RFI, RFM, RFSa, RFSb, RPI, RPSa, RPSb, RTB | RFSb1, RFSb2 |
| P43 | MGH | 7 | ANT, PuO, PuM, Unknown, VL | 13 | 7L+7R | 50.17 | LAMY, LFIa, LFIb, LHH, LHT, LTSa, LTSb, RAMY, RFIa, RFIb, RHH, RHT, RTSa, RTSb | LAMY1, LAMY2, LHH1, LHH2, LHH3, RAMY1, RAMY2, RHH1, RHH2, RHH3 |
| P44 | MGH | 6 | CM, MD, PuO, PuM, VL | 9 | 7L+1R | 30.44 | FSS, LFSI, LHH, LPI, LPP, LPS, LTB, LTS, RPS | LFSI11, LFSI12, LFSI13, LFSI14, LFSI15, LFSI16 |
| P45 | MGH | 4 | PuM | 4 | 9R | 59.5 | RAMY, RFI, RHH, RHT, RPI, RTB, RTO, RTSa, RTSb | RAMY1, RAMY2, RAMY3, RHH1, RHH2, RHH3, RHH4 |
| P46 | MGH | 3 | PuO, PuM, VL | 19 | 15L+6R | 31 | LAMY, LFIa, LFIb, LFS, LHH, LHT, LO, LPC, LPI, LPOc, LPOi, LPOs, LPS, LTB, LTS, RFIa, RFIb, RO, RPOs, RT, RTB | LPOc1, LPOc10, LPOc2, LPOc3, LPOc4, LPOc5, LPOc6, LPOc7, LPOc8, LPOc9, LPOi13, LPOi14, LPOi15, LPOi16, LTS11, LTS12, LTS13, LTS14, LTS15, LTS16, RPOs1, RPOs2, RPOs3, RPOs4 |
| P47 | MGH | 6 | CM, VL | 14 | 6L+7R | 31 | LAMY, LFI, LFMa, LFMb, LFMc, LHH, RAMY, RFI, RFMa, RFMb, RFMc, RFS, RHH | LFI4, LFI5, LFI6, RAMY1, RAMY2, RAMY3, RFI10, RFI11, RFI12, RFI13, RFI14, RFI9, RFMa10, RFMa8, RFMa9, RFMb1, RFMb2, RFMb3, RFMb4, RFMb5, RFMb6, RFMc12, RFMc13, RFMc14, RFMc15, RFMc16, RFS6, RFS7 |

| | | | | | | | | |
|---|---|---|---|---|---|---|---|---|
| P48 | MGH | 8 | PuO, PuM, VL | 9 | 8L+4R | 31 | LAMY, LHH, LHT, LO, LSMa, LTB, LTM, LTO, RHH, RHT, RTB, RTM | LHH1, LHH10, LHH11, LHH12, LHH13, LHH14, LHH2, LHH3, LHH4, LHH5, LHH6, LHH7, LHH8, LHH9, LHT1, LHT10, LHT11, LHT12, LHT13, LHT14, LHT15, LHT16, LHT2, LHT3, LHT4, LHT5, LHT6, LHT7, LHT8, LHT9, LTB1, LTB10, LTB11, LTB12, LTB13, LTB14, LTB15, LTB16, LTB2, LTB3, LTB4, LTB5, LTB6, LTB7, LTB8, LTB9, LTM10, LTM11, LTM12, LTM13, LTM14, LTM15, LTM16, LTO4, LTO5, LTO6, LTO7, LTO8 |
| P49 | MGH | 3 | CM, PuO, PuM, VL | 10 | 11L+3R | 31 | LAMY, LFI, LHH, LHT, LPIa, LPIb, LPIc, LPId, LPS, LTSa, LTSb, RAMY, RHH, RTS | LFI10, LFI11, LFI12, LFI13, LFI14, LPIb1, LPIb2, LPIb3, LPIb4, LPIb5, LPIb6, LPId1, LPId2, LPId3, LPId4, LPId5, LPId6, LPId7, LTSb1, LTSb2, LTSb3, LTSb4, LTSb5, LTSb6 |
| P50 | MGH | 5 | CM, MD, VL | 14 | 5L+6R | 31 | LFIa, LFIb-A, LFMa-D, LFMb-C, LFS, RFIa, RFIb-A, RFMa, RFMb-C, RFSa-D, RFSb-M | RFIa10, RFIa11, RFIa12, RFIa13, RFIa14, RFMa10, RFMa11, RFMa12, RFMa13, RFMa14, RFMb-C1, RFMb-C2, RFMb-C3, RFSa-D5, RFSa-D6, RFSa-D7, RFSa-D8, RFSa-D9 |
| P51 | MGH | 4 | ANT, CM, PuM, VL | 7 | 3L+8R | 31 | LAMY, LHH, LHT, RAMY, RFIa, RFIb, RFIc-A, RFMa-C, RFMb, RHH, RHT | RAMY1, RAMY2, RAMY3, RAMY4, RFIa1, RFIa2, RFIa3, RFIb1, RFIb2, RFIb3, RFIb4, RFMb1, RFMb2, RFMb3, RFMb4, RFMb5, RFMb6, RFMb7, RHH1, RHH2, RHH3, RHH4, RHT1, RHT2, RHT3, RHT4 |
| P52 | MGH | 10 | ANT, CM, VL | 11 | 2L+9R | 27.49 | LFO-CM, LFS, RAMY, RFMa, RFMb-A, RFO-CM, RFSa, RFSb, RFSc, RFSd, RPI | RFSb10, RFSb8, RFSb9, RFSc10, RFSc8, RFSc9 |
| P53 | MGH | 6 | CM, MD, VL | 7 | 14L | 31 | LFIa-A, LFIb, LFIc-C, LFId, LFMa, LFMb, LFMc-P, LFSa, LFSb-S, LFSc, LFSd, LFSe, LPS, LTM | LFId1, LFId2, LFId3, LFMA1, LFMA2, LFMA3 |
| P54 | MGH | 11 | ANT, MD, PuO, PuM, VL | 13 | 8L+8R | 30.27 | LAMY, LFI, LHH, LHT, LTB, LTSa-A, LTSb, LTSc-P, RAMY, RFIa, RHH, RHT, RTB, RTSa-A, RTSb, RTSc-P | LAMY1, LAMY2, LAMY3, LAMY4, LAMY5, LAMY6, LHH1, LHH2, LHH3, LHH4, LHH5, LHT1, LHT2, LHT3, LHT4, RAMY1, RAMY2, RAMY3, RH1, RH2, RH3, RHH1, RHH2, RHH3, RHH4 |
| P55 | MGH | 7 | ANT, Unknown, VL | 6 | 8L+8R | 55.71 | LAMY, LFIa, LFIb, LHH, LHT, LTB, LTP, LTS, RAMY, RFIa, RFIb, RHH, RHT, RTB, RTP, RTS | LHH1, LHH2, LHH3, LHH4, LHH5, LHT1, LHT2, LHT3, LHT4, LHT5, RHH1, RHH2, RHH3, RHH4, RHH5, RHT1, RHT2, RHT3, RHT4, RHT5 |

##### Table S2: Long-timescale LMM test statistics

| Effect | F | β | DF1 | DF2 | p |
|---|---|---|---|---|---|
| **Spindle rate ~ IED rate × Day + (1\|PatientHemiID)** | | | | | |
| Intercept | 218.13 | 7.2312 | 1 | 448 | **<0.001** |
| Day | 27.259 | -0.53969 | 1 | 448 | **<0.001** |
| IED rate | 8.2491 | -0.25532 | 1 | 448 | **0.0043** |
| Day × IED rate | 2.1021 | -0.13218 | 1 | 448 | 0.1478 |

Spindle rate ~IED rate × Day + (1|PatientHemi)

##### Table S3: Short-timescale LMM variable descriptions

| Effect | Variable definition |
|---|---|
| Y | Outcome variable; either suppression index or facilitation index |
| IED rate | Z-scored SOZ-IED rate; continuous variable |
| Spindle rate | Z-scored thalamic spindle rate; continuous variable |
| NucleusID | Thalamic nucleus category: ANT, CM, MD, PuM, PulO, and VL |
| TC | Binary variable: 0/1; whether the pair has significant thalamus → SOZ Granger causality |
| CT | Binary variable: 0/1; whether the pair has significant SOZ → thalamus Granger causality |
| f | Frequency-band category: 11-12 Hz, 12-13 Hz, 13-14 Hz and 14-15 Hz |
| Δt | Z-scored thalamic spindle duration; continuous variable |

*Abbreviations: SOZ, seizure onset zone; IED, interictal epileptiform discharge; VL, ventrolateral; PuM, medial pulvinar; PuO, other pulvinar; CM, centromedian; MD, medial dorsal; ANT; anterior nucleus of the thalamus.*

##### Table S4: Short-timescale LMM test statistics

| Y | Effect | F | β | DF1 | DF2 | p |
|---|---|---|---|---|---|---|
| | **Y ~ 1 + NucleusID × (TC + CT) + f + Δt + IED rate + Spindle rate + RE** | | | | | |
| **SI** | Intercept | 13.22 | 0.0019424 | 1 | 24639 | **<0.001** |
| **FI** | | 24.439 | 0.0028671 | 1 | 24639 | **<0.001** |
| **SI** | NucleusID | 0.75619 | — | 5 | 24639 | 0.58137 |
| **FI** | | 2.8221 | — | 5 | 24639 | **0.014943** |
| **SI** | TC | 0.59501 | 0.00014674 | 1 | 24639 | 0.4405 |
| **FI** | | 1.7086 | 0.00025612 | 1 | 24639 | 0.19118 |
| **SI** | CT | 0.14039 | 0.00007488 | 1 | 24639 | 0.70789 |
| **FI** | | 16.094 | 0.00082692 | 1 | 24639 | **<0.001** |
| **SI** | NucleusID × TC | 4.2055 | — | 5 | 24639 | **<0.001** |
| **FI** | | 1.8008 | — | 5 | 24639 | 0.10894 |
| **SI** | NucleusID × CT | 1.0099 | — | 5 | 24639 | 0.4099 |
| **FI** | | 3.8421 | — | 5 | 24639 | **0.0017609** |
| **SI** | f | 9.5185 | — | 3 | 24639 | **<0.001** |
| **FI** | | 11.923 | —- | 3 | 24639 | **<0.001** |
| **SI** | Δt | 0.03988 | 0.00002868 | 1 | 24639 | 0.84172 |
| **FI** | | 18.127 | 0.00061817 | 1 | 24639 | **<0.001** |
| **SI** | IED rate | 290.03 | 0.0022777 | 1 | 24639 | **<0.001** |
| **FI** | | 89.393 | -0.0013646 | 1 | 24639 | **<0.001** |
| **SI** | Spindle rate | 22.187 | -0.0008014 | 1 | 24639 | **<0.001** |
| **FI** | | 43.7 | 0.0011296 | 1 | 24639 | **<0.001** |

RE = (1|PatientHemiID) + (1|PatientHemiNightID) + (1|SOZChannelID) + (1|ThalamusChannelID)

* Since TC and CT are binary variables, $\beta$ represents the difference between 1 and 0 (presence – absence) of directed connectivity
